# Genome profiles of lymphovascular breast cancer cells reveal multiple clonally differentiated outcomes with multi-regional LCM and G&T-seq

**DOI:** 10.1101/807156

**Authors:** Zhongyi Zhu, Weiwei Wang, Feng Lin, Tracy Jordan, Guibo Li, Sveta Silverman, Si Qiu, Anil Abraham Joy, Chao Chen, Deanna L. Hockley, Xi Zhang, Qing Zhou, Lynne M. Postovit, Xiuqing Zhang, Yong Hou, John R. Mackey, Bo Li, Gane Ka-Shu Wong

## Abstract

Lymphovascular invasion (LVI) is a critical step in the metastatic process but have received relatively little attention due to the technical challenges associated with their isolation. In this study, we used laser capture microdissection (LCM) to isolate 97 cancer cell clusters from pathological frozen sections within lymphatic vessels, primary tumor tissue, and axillary lymph nodes of a triple negative breast cancer (TNBC) patient. Simultaneous genome and transcriptome amplification and sequencing (G&T-seq) performed on these clusters permitted a comprehensive depiction of the genomic and transcriptional profiles of cancer cells associated with LVI. Combination phylogeny analysis pointed to three evolutionarily distinct pathways of tumor clone development and metastasis in this patient, each of which was associated with a unique mRNA signature, and correlated to disparate overall survival outcomes. Moreover, hub gene evaluation found extensive down regulation of ribosomal protein mRNA to be a potential marker of poor prognosis in breast cancer patients.

## INTRODUCTION

Tumor metastasis is a highly complex process that occurs through several steps, including tumor cell invasion of local tissue, intravasation and survival of tumor cells in the circulation, arrest and extravasation at a distant organ, micrometastasis formation, and finally metastatic colonization^1^. Cancer cells can enter the circulation either directly by invading tumor associated blood vessels (termed blood vessel invasion or BVI), or indirectly by first traversing the lymphatic system in a process known as lymphovascular invasion (LVI)^2,3,4^. In breast cancer, the majority of vessel intravasion takes place at the primary tumor location via LVI and the lymphatic network serves as a conduit for the metastatic spread of this disease^2, 5, 6^. Once within the lymphatic vasculature, breast cancer cells are diverted, along with immune cells and lymphatic fluid, to the local regional axillary lymph nodes where their outcome can vary from indefinite lymph node arrest, to entry into the circulation and subsequent formation of distant metastatic lesions^4^. The rate at which cancer cells enter the lymphatic system and transit to the lymph nodes is influenced by a number of factors, including biomechanical forces from within the primary tumor and the lymphatic vessels, and the presence of chemokines and other immune elements that alter the primary tumor and lymphatic microenvironment^2, 4, 7,8,9^. Because the majority of breast cancer deaths result from metastatic spread^10^, isolating and characterizing metastatic invasive cancer cells is often pursued as way to identify new medicinal targets to prevent or impede metastasis in breast cancer patients.

Tumorigenesis itself occurs by way of genetic aberrations that alter function and/or expression of specific “driver genes”, or inactivate tumor suppressors. For disease progression to occur, cancer cells are believed to acquire further mutations which promote expansion, invasion of surrounding tissues, and finally, tumor metastasis^11^. Selection pressure from the local microenvironment and/or systemic drug treatment can aid this process by promoting the expansion of tumor cell clones harboring common genetic or transcriptional abnormalities advantageous to these cells^11,12,13^. Thus, at any time point in the tumor lifecycle, there exists the possibility of detecting multiple tumor cell clones within a single patient^14^. Clonal populations contribute to intra-tumor heterogeneity and are of particular relevance as they have the potential to represent drug-resistant populations, or cells with a predisposition for metastatic spread^15^. The detection of clonal populations has been greatly advanced by the advent of single cell sequencing which has facilitated the evaluation of genetic mutations throughout the genome of a single cell. This has allowed us to attribute unique DNA and RNA signatures to tumor cells based on their presence in specific microenvironments. In the case of breast cancer, single cell sequencing has helped define gene expression signatures related to metastatic burden^16^, metastatic subtype^17^, and even spatial orientation within primary breast cancer tissue^18, 19^. However, methods for isolating single cells, as well as subsequent DNA and RNA sequencing steps, vary widely and few groups have analyzed both genomic aberrations and gene expression changes together in a particular cancer cell type. Furthermore, sequencing single cells may not be the ideal method for characterizing metastatic tumor populations as tumor cells that circulate as clusters exhibit higher metastatic potential, and are associated with worse disease prognosis^20, 21^. These circulating tumor cell clusters are often also polyclonal, demonstrating a mix of primary tumor and epithelial-like characteristics that are believed to enhance their invasiveness, dissemination, and metastatic colonization^20,21,22^. Thus, analyzing invasive tumor cells as a collective, multicellular unit rather than on a single-cell level, may provide a more physiological representation of cancer cells with an aggressively metastatic phenotype.

In this report, we isolated and characterized clusters of breast cancer cells from the tissues of a triple negative breast cancer (TNBC) patient using a combination of laser-capture microdissection (LCM) and G&T-seq^23^. This combination of techniques allowed us to not only extract and characterize clusters of tumor cells from the primary and lymph node tumors of this patient, but also directly from within lymphatic vessels, thereby enabling the analysis of cells discernably *en route* to the axillary lymph nodes from the primary tumor. G&T-seq, which permits segregation and sequencing of genomic DNA and full-length mRNA transcripts simultaneously from a single cell, was then performed on these cell clusters in parallel to assess the chromosome aberration patterns, genetic mutations, and gene expression profiles associated with LVI and lymph node metastasis. By this method, we identified a genomic aberration pattern and RNA expression profile associated with LVI. Moreover, we found evidence of three transcriptionally distinct pathways of metastatic spread within a single patient which, when extrapolated to a larger patient population, were found to be varying predictors of breast cancer survival outcomes. Thus, the combined utilization of these techniques may help identify new targets for preventing metastatic disease in cancer patients, and provide a valuable tool in the development of personalized medicine for patients with known heterogeneous diseases like TNBC.

## RESULTS

### Cell cluster isolation from primary breast tumor and axillary lymph nodes using LCM

To isolate and characterize cancer cells with an inherent ability to invade the lymphovasculature and form distant metastatic lesions, we obtained fresh tissue samples from a 24 year old woman with T3 N1 M1, ER(-), PR(-), HER-2(-), grade 3, invasive ductal breast carcinoma following palliative mastectomy and full axillary lymph node dissection. This patient had treatment refractory disease (two cycles of docetaxel and four cycles of doxorubicin/cyclophosphamide chemotherapy) which, at the time of sample collection, had metastasized to the patient’s local regional axillary lymph nodes and liver. To maximize the likelihood of detecting LVI, we collected two tissue samples from the tumor-stromal interface of the primary tumor, as well as two samples from her carcinoma positive axillary lymph nodes (3 of the 15 removed lymph nodes were positive for metastatic disease). These tissue samples were flash-frozen, sectioned, H&E-stained, and screened by an anatomic pathologist for the presence of cancer cell clusters defined as histologically malignant groups of cells of at least 10µm in diameter^21, 24^; Fig.1). Laser capture microdissection (LCM), which permits the acquisition of select cells while preserving anatomical structures such as lymphatic vessels, was then utilized to dissect cell clusters from select tissue sections. A total of 186 cell clusters were collected for processing with 97 (∼52%) passing both DNA and RNA quality control checks for sequencing analysis (Table S1.1). These clusters were comprised of 17 morphologically normal breast epithelial cell clusters, 17 primary tumor cancer cell clusters, and 20 clusters found within lymphatic vessels which, due to the unidirectionality of lymphatic flow, represent cells categorically *en route* to the axillary lymph nodes from the primary tumor. Additionally, 36 cancer cell clusters and 5 lymphocyte cell clusters were collected from the tumor-infiltrated axillary lymph node samples. G&T-seq was then performed as described ^23^ to simultaneously extract, amplify, sequence, and create DNA and mRNA libraries from each LCM isolated cell cluster. In brief, multiple-displacement amplification (MDA) was performed on the genomic DNA (gDNA) isolated from each cluster, followed by whole genome (WGS) and whole exome sequencing (WES) to detect copy number variants (CNV) and single nucleotide variants (SNV), respectively. Whole genome sequencing (WGS) was performed at a sequencing depth of 1.12x while whole exome sequencing (WES) was performed at a usage depth of 272X (Raw depth >1000X). Whole transcriptome sequencing (WTS) was performed on mRNA isolated from each cluster at a sequencing depth of 4.5G clean base with a quality score of 30 or higher.

**Figure 1.**
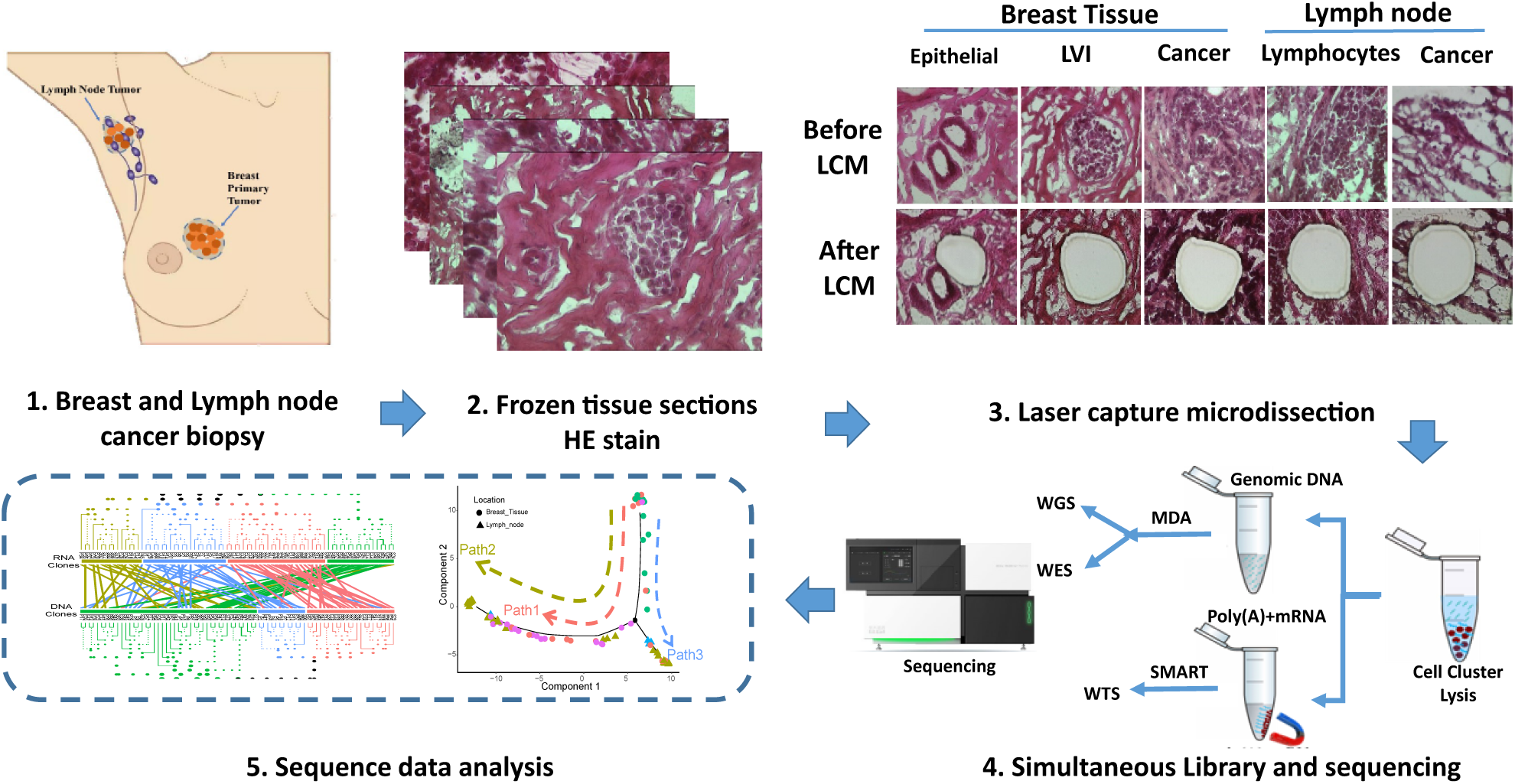
Schematic representation of LCM cell cluster isolation and G&T-seq from TNBC patient samples. Tissue samples were collected from the tumor-stromal interface of the primary tumor, and carcinoma positive axillary lymph nodes of a single TNBC patient. Tissue samples were flash-frozen, sectioned, and H&E stained for pathological examination. LCM was used to isolate epithelial, LVI, lymphocyte and cancer cell clusters measuring >10µm in size from anatomically intact tissue sections. G&T-seq was then performed to extract and sequence gDNA and mRNA from each cell cluster. gDNA was amplified with MDA, and mRNA was amplified using a modified Smart-seq2 protocol. WGS and WES was performed on the amplified gDNA for CNV and SNV detection, respectively, while WTS was performed on amplified mRNA transcripts for transcriptome analysis.

### LVI associated cell clusters share a CNV profile abundant in chromosome amplification and deletion events

To begin our assessment of the LCM isolated cell clusters, we performed WGS on the gDNA isolated from each cluster to detect the presence of CNVs. Additionally, we sought to determine if a relationship existed between the cell types sampled, and the CNV pattern identified amongst individual cell clusters, consistent with the existence of CNV-defined clonal populations. While historically the term “clone” has been used to define genetically or transcriptionally identical cells, the cell clusters assessed in these experiments were inherently heterogeneous. As such, we defined “clones” as cell clusters sharing a common genomic or transcriptional profile. Using the single cell CNV calling method previously described by Baslan *et al* to avoid amplification bias^25^, we detected chromosome amplification and deletion events present in every cell cluster analyzed (Fig.S1). This included clusters comprised of histologically normal cells positioned in a well-organized breast duct, indicative of the heterogeneous nature of the cell clusters isolated from this patient. Next, we performed hierarchical clustering using the Ward.D2 algorithm to generate a CNV heatmap to evaluate the clonal architecture of these clusters, using bootstrap to identify stable clustering clades. Heatmap analysis showed the existence of three distinct CNV profiles which we appointed CNV Clone A, CNV Clone B and CNV Clone C (Fig.2A). Notably, cell clusters obtained from within lymphatic vessels clustered together as CNV Clone B (CNV LVI clone, pink) along with several primary cancer cell clusters. Therefore, not only did the LVI associated cell clusters exhibit a specific CNV profile, this profile was shared with a number of primary cancer cells. By comparison, cell clusters associated with CNV clone C (CNV LN clone, blue) were isolated exclusively from the lymph nodes of this patient. Because no cell type heterogeneity was associated with CNV clone C, this indicates that we either didn’t sample the closest ancestor of this lymph node clone in primary tumor, or the phenotype of this ancestor changed significantly during the course of disease progression. Lastly, CNV Clone A (CNV mixed-epithelial clone, green) was found to be comprised by a combination of epithelial, lymphocyte, primary cancer and lymph node cancer cell clusters. These clusters exhibited relatively few copy number variations (4.1%), but shared a common amplification of the X chromosome. Notably, because the cell clusters associated with the mixed-epithelial CNV clone included lymph node cancer cells, this suggests that relatively few CNV changes were required to promote lymph node metastasis in this patient. The LVI and LN CNV clones by contrast exhibited a substantial number of CNV changes, characteristic of chromosome instability (CIN). The LVI clone (CNV clone B) contained CNVs which encompassed approximately 71.7% of the entire genome, and included amplification of the X chromosome. The LN clone (CNV clone C) also exhibited extensive CNV changes covering ∼53.1% of the genome, but these did not include a significant X chromosome amplification (Fig.2A, Table S2.1). In summary, these results indicate the presence of a unique, genomically unstable LVI clone in this patient, as well as two CNV-defined metastatic lymph node clones associated distinct amounts of CIN.

**Figure 2.**
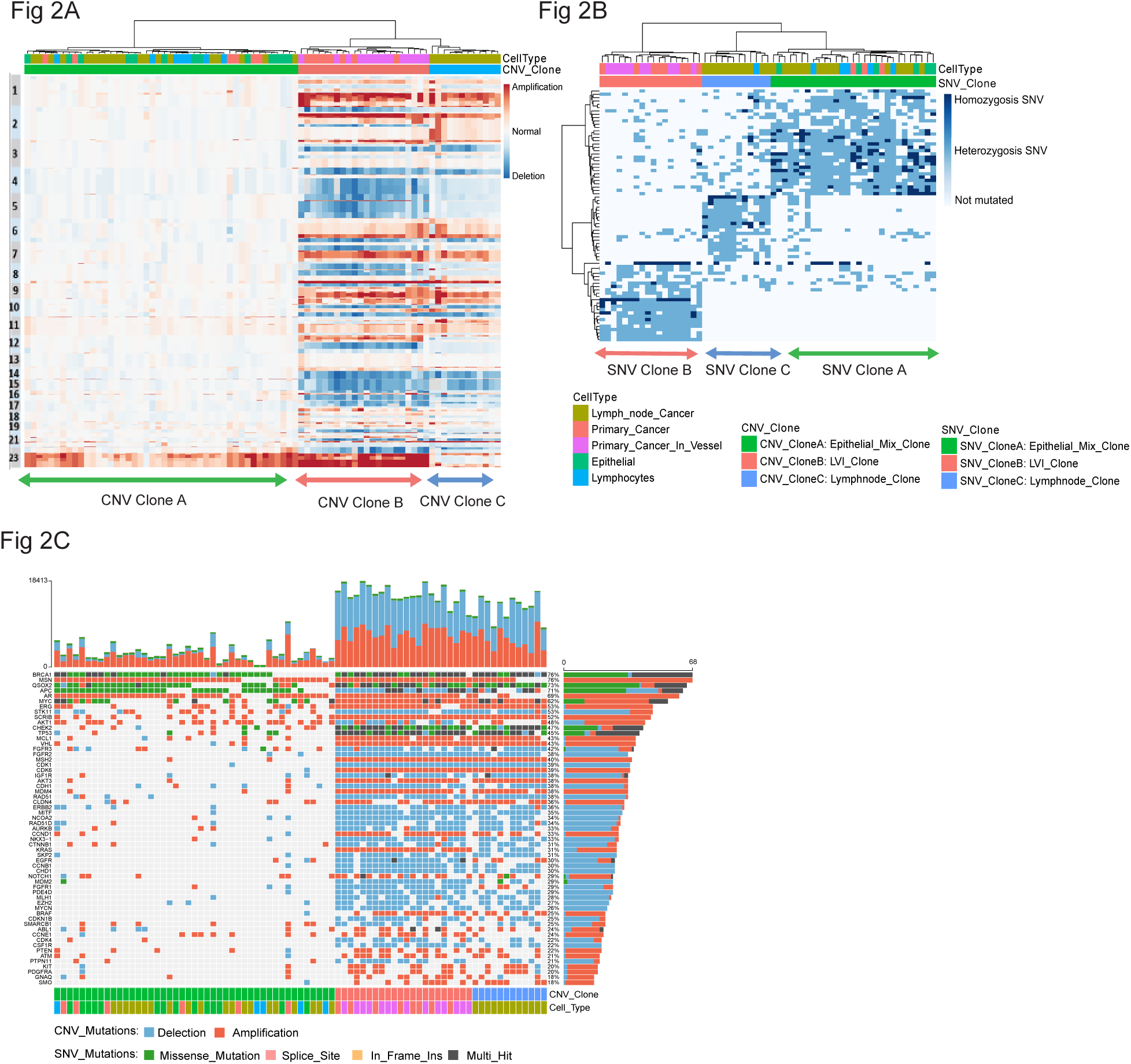
CNV and SNV sequencing reveals the presence of a distinct LVI cancer cell clone amongst a vast mutational landscape. (A) CNV heatmap generated by unsupervised hierarchal clustering of 80 cell clusters following WGS. Cell clusters are plotted along the x-axis, and CNVs are plotted in genomic order along the y-axis. Color bars (top) represent cell type (yellow, LN; red, primary tumor; pink, primary tumor associated vessels; green, epithelial; blue, lymphocytes), and clone designation (green, mixed clone; blue, LN clone; pink, intravasation clone). Chromosomal amplifications are depicted in red while deletions are shown in blue. (B) SNV heatmap generated by unsupervised hierarchal clustering of 59 cell clusters following WES. Cell clusters are plotted along the x-axis, and SNVs are plotted according to clone prevalence along the y-axis. Cell type and clone designation is indicated as in (A). Homozygous SNVs are depicted as dark blue, heterozygous SNVs are light blue, and genes with no mutations are white. (C) Oncoplot depicting chromosome alterations and/or genetic mutations detected in 59 common oncogenes. Each vertical column represents the CNV (amplification, red; deletion, blue), SNV (missence, green; splice site, pink; frame shift, yellow) or multi-hit (black) mutations detected in each oncogene from the gDNA of each cell cluster following WGS and WES sequencing. Stacked bars (top) show the accumulated alterations (both CNV and SNV) across all 23,588 ref genes in each cell cluster. Stacked bars (right) show the accumulated number of CNV and SNV mutations detected in each oncogene across all 80 cell clusters, and the percentage of cell clusters harboring mutations in each oncogene. Individual cell clusters are arranged based on CNV clone association and oncogenes are arranged by mutation/chromosome alteration frequency.

### Cell clusters associated with LVI have a high homozygous mutational burden

Next, we performed WES on the gDNA isolated from each LCM cell cluster to detect the presence of SNVs, and to verify the existence of potential SNV clones based upon common gene mutation patterns. We again performed unsupervised clustering of each LCM cluster and used the data generated to construct SNV heatmaps whereby the homozygous and heterozygous SNVs detected in each cell cluster were depicted^26^. As represented by Fig.S2 in which all SNVs detected are illustrated, 1704 clonal specific SNVs were identified in the tumor samples from this patient. Similar to our CNV findings, clustering analysis of the SNVs identified showed the existence of three distinct SNV clones based upon shared gene mutation patterns (Fig.2B). SNV clone A (mixed-epithelial clone, green) was comprised of LCM cell clusters representing every cell type tested with the exception of lymphatic vessel associated cell clusters, and harbored SNVs occurring at a range of frequencies (6.7% - 100%) in over 750 genes (Table S2.2). By contrast, SNV clone B contained cell clusters isolated from the primary tumor and lymphatic vessels of this patient, again indicating the presence of an LVI clone (SNV clone B, pink). Finally, a number of LCM cell clusters isolated from the lymph nodes exhibited a similar gene mutation pattern indicative of a SNV lymph node clone (SNV clone C, blue). Mutations were detected in both the LVI and LN SNV clones at a range of frequencies (LVI SNV clone 11.1% - 100%, LN SNV clone 18.2% - 100%), affecting over 540 and 450 genes, respectively (Table S2.2). Furthermore, cell clusters associated with the SNV LVI clone were found to have, on average, a higher homozygous mutation burden than other cell clusters (Fig.S2), suggesting that the degree of SNV homozygosity may be related to LVI invasiveness.

### Lymphovascular-invasive cancer cells contain numerous oncogenic mutations including *BRCA1* and *TP53*

Utilizing our WGS and WES results, we next sought to identify any potentially significant oncogenic events which may have occurred in this patient. We compiled a list of over 89 known transformation associated genes and assessed each cell cluster for the frequency of SNV missense, splice site, and frame shift mutations, as well as CNV associated chromosome deletion and amplification events in these genes. In total, 59 of these genes were found to harbor either SNV or CNV mutations (Fig. 2C). As commonly observed in young TNBC patients ^27,28,29^, the DNA repair associated gene *BRCA1* contained either a missense mutation, was deleted, or exhibited multiple transformative hits in ∼76% of all of the cell clusters isolated, including those of mixed-epithelial origin (CNV clone A; Fig.2C). Other genes affected by single nucleotide variability included the tumor suppressor *TP53*, which was either mutated or exhibited multiple hits in the LVI (CNV clone B) and LN (CNV clone C) associated clones, and *APC* which was mutated in ∼71% of all the cell clusters. In accordance with an amplification of the X chromosome, genes *AR* and *MSN* located on the X chromosome were amplified in a high proportion of mixed-epithelial (CNV clone A) and LVI (CNV clone B) CNV clone associated cell clusters, but to a lesser degree in the LN CNV clone (CNV clone C) (Table S2.3). Furthermore, *MYC, ERG, MCL1, VHL, MSH2* and *CDK6* were found to be amplified while *STK11, FGFR2, IGF1R* were deleted in nearly all of the LVI (CNV clone B, pink) and LN (CNV clone C, blue) associated cell clusters. Overall this demonstrates the complex CNV and SNV landscape in this patient with genetic alterations affecting not only the CIN associated LVI and LN clones, but also the mixed-epithelial CNV/SNV clones.

### Two transcriptional distinct lymph node RNA clones identified using whole transcriptome sequencing

In metastatic breast cancer, the tumor microenvironment varies considerably among the primary tumor, the tumor associated lymphatic vessels, and the sentinel lymph nodes. As such, we aimed to determine if the isolated cell clusters had common gene expression patterns characteristic of RNA-defined tumor cell clones. WTS was performed on the mRNA extracted from each LCM cell cluster to detect the relative expression of 23,588 RefSeq curated genes. Following mRNA library construction, we used the algorithm SC3 to perform unsupervised clustering of each cell cluster based upon their gene expression pattern, and generated a RNA heatmap whereby individual cell clusters are grouped together based upon their relative expression of select marker genes^30^ (Fig.3A). By this method, we detected four RNA clones within the tissue samples of this patient (Fig.3A), each of which was confirmed by silhouette plots, and by calculating the stability index for each clade. The first clone identified was termed a RNA epithelial clone (RNA clone A, green) as cell clusters with this gene expression pattern were predominantly histologically normal breast epithelial cells. Again, LCM cell clusters isolated from lymphatic vessels grouped together and were found to exhibit a unique RNA expression pattern, indicating the presence of a RNA LVI clone (RNA clone B, pink). Marker genes highly expressed by cell clusters with an LVI clone phenotype included several related to tumor invasion and cell differentiation including *FOXC1/D1/Q1, NOTCH1, ART3, BIRC7, RAB40B, PTP4A3, CDK1, CLDN4/7, FGFR3, QSOX2, AURKB, SCRIB* and *CCNB1* (Table S3.1). Unlike our CNV and SNV analysis, transcriptional assessment detected the presence of two distinct lymph node RNA clones, each with a unique pattern of gene expression. RNA LN clone 2 (RNA clone D, yellow) exhibited very high expression of a number of immune-related marker genes such as *CD163, HMOX1* and *TYROBP*, as well as several involved with cell migration including *VCAN* and *CTSH* (Table S3.1). RNA LN clone 1 (RNA clone C, blue) by comparison demonstrated a relatively muted level of gene expression with *POMK*, *CYP3A4*, and several solute carriers being amongst the most highly expressed markers genes (Table S3.1). Thus, while these lymph node clones are related in terms of their physical location, their gene expression profiles are distinct suggesting they are unrelated genomically.

**Figure 3.**
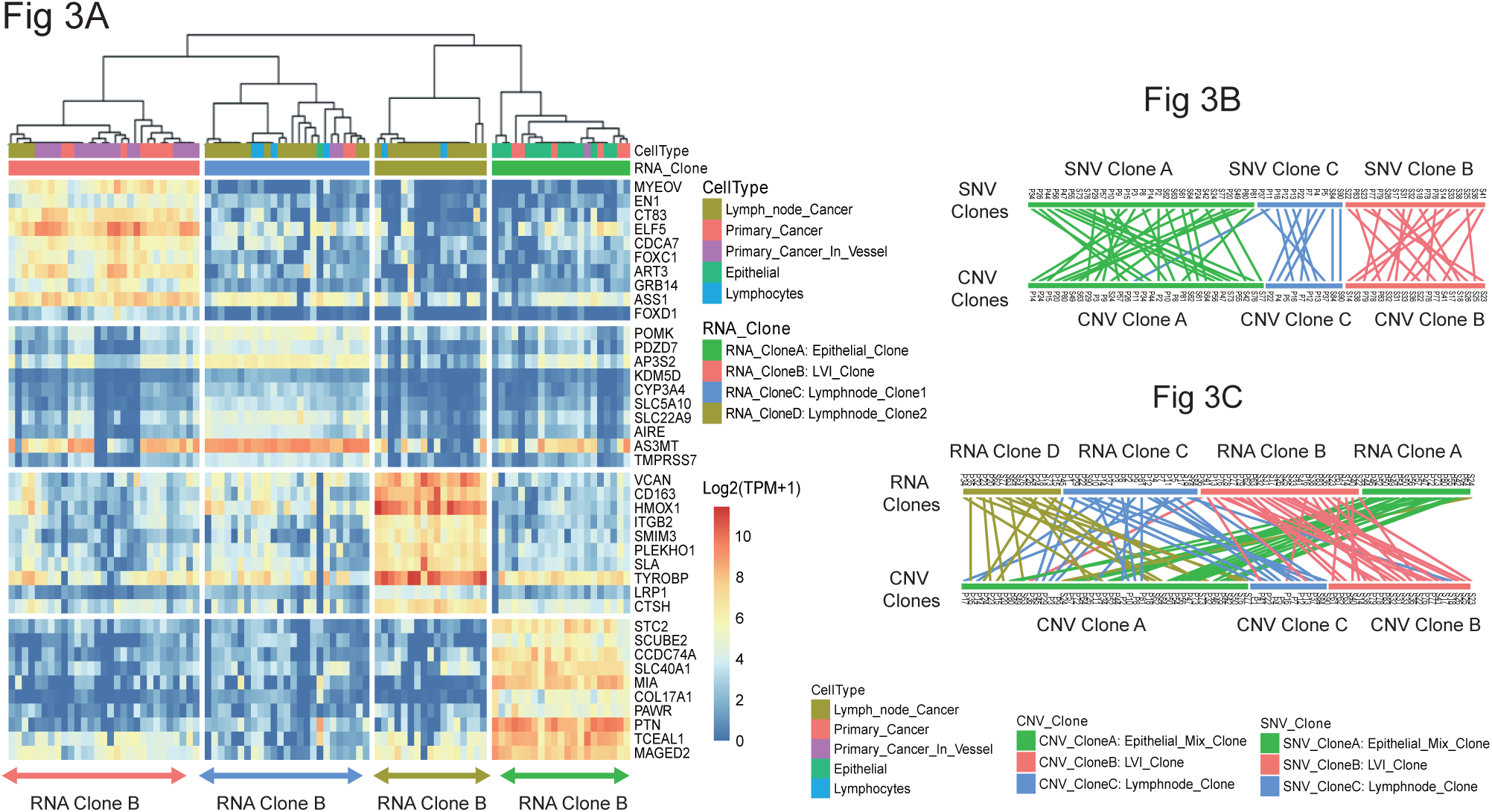
G&T-seq identifies four RNA-based clonal population in a single TNBC patient and permits direct comparison between DNA and RNA clones. (A) RNA heatmap depicting the unsupervised hierarchal clustering of 92 cell clusters based on their relative expression of select marker genes. Cell clusters are plotted along the x-axis and marker genes are plotted along the y-axis. Color bars (top) represent the cell type (yellow, LN; red, primary tumor; pink, primary tumor associated lymphatic vessels; green, epithelial cells; blue, lymphocytes) and RNA clone designation (green, epithelial clone; blue, LN subclone 1; yellow, LN subclone 2; pink, LVI clone). Increases in relative gene expression are represented on a Log2 (TPM+1) scale where 10 (light red) is equivalent to >1000-fold increase and 0 (dark blue) indicates no relative increase in gene expression. (B) Tanglegram comparison of SNV clones (top) and CNV clones (bottom). Clonal subpopulations are depicted by color (green, mixed; blue, LN; pink, intravasation). (C) Tanglegram comparison of RNA clones (top) and CNV clones (bottom). Clone are depicted by color (RNA: green, epithelial; pink, intravasation; blue, LN subclone 2; yellow, LN subclone 1; CNV: green, mixed; blue, LN; pink, intravasation). Auxiliary lines in tanglegrams connect corresponding LCM cell clusters.

### G & T sequencing enables direct comparison between DNA and RNA clones

Because G&T-seq allowed us to extract and analyze DNA and RNA from the same tissue samples, we sought to validate the clonal assignment of our cell clusters and confirm the relationships among the CNV, SNV, and RNA clones identified. To visualize these relationships, we employed tanglegrams generated using the R package software Dendextend^31^, whereby two phylogenetic trees are drawn opposite of each other, and auxiliary lines are used to connect corresponding cell clusters^32, 33^. Tanglegrams comparing the relationship between the CNV and SNV clones demonstrated that, with the exception of a single cell cluster, cell clusters designated as a CNV A, CNV B, or CNV C clone mapped to a corresponding SNV A, SNV B or SNV C clone (Fig.3B). This indicates that on a DNA level, the CNV and SNV clones identified are relatively stable and the cell clusters associated with these clones exhibit a unique CNV and SNV profile. Using the same technique, we then compared the relationship between CNV and RNA clones (Fig.3C). While not as categorically definitive as our DNA results, we found a significant amount of overlap between clones despite having identified three CNV clones and four RNA clones. Specifically, 84% of the LCM cell clusters with a LVI RNA clone phenotype (RNA clone B, pink) corresponded to a matching LVI CNV clone (CNV clone B, pink), and 100% of the cell clusters with a RNA epithelial clone phenotype (RNA clone A, green) corresponded to a matching mixed-epithelial CNV clone (CNV clone A, green). As expected however, more disorder was observed when comparing LN clones. While 100% of the LCM cell clusters with a RNA LN clone 2 phenotype (RNA clone D, yellow) corresponded with the mixed-epithelial CNV clone (CNV clone A, green) exhibiting relatively few CNVs, cell clusters associated with RNA LN clone 1 (RNA clone C, blue) corresponded with both the mixed-epithelial (CNV clone A, green) and LN CNV clone (CNV clone C, blue), previously found to have a high number of CNVs. Hence, this analysis not only indicated that the CNV/SNV clones are largely comparable to their RNA clone counterparts, it also confirmed that the two RNA lymph node clones are transcriptionally distinct due to difference in chromosome copy number.

In circumstances where individual cell clusters matched well to a CNV, SNV and RNA clone, we also generated Venn diagrams to depict commonalities between genes affected by CNV, SNV, and expression changes (Fig.S3). Little mutual overlap was observed in genes associated with either the mixed-epithelial clone or LN clone 2 indicating that few genetic alterations exist that can be used to predict the phenotype of these clones. However, the number of overlapping genes, particularly those affected by both CNV and SNV changes, was higher in cell clusters associated with LN clone 1 and the LVI clone (Fig.S3). This is consistent with these clones exhibiting a large number of genomic alterations, and the LVI clone being the most homogeneous in terms of cell type composition. Furthermore, the number of overlapping genes between CNV and RNA clones was general much higher compared to SNV and RNA clones. This suggests that while point mutations may have played a role in shaping the transcriptome of the clones isolated to some degree, CNVs appeared to have a larger influence on transcription in this patient.

### G&T-seq conveys the evolutionary history of tumor cell clones as well as pathways of metastatic spread in this patient

Having established the relationships among the LVI, lymph node, and mixed/epithelial clones, we were interested in determining how these clones were evolutionarily related. To this end, we constructed a CNV maximum parsimony tree that allowed us to trace the existence of common ancestors, and measure the evolutionary distance between each CNV clone^18, 26, 34^. LCM cell clusters were plotted against hamming distance and further categorized based upon cell type and tissue sample location (lymph node vs. breast tissue). Overall, the CNV phylogenetic tree was rooted by the mixed-epithelial CNV clone (CNV clone A, green) with relatively few CNVs, and showed evidence of both branched and gradual evolution (Fig.4A). Two early divergent subpopulations were identified branching from the mixed epithelial clone representing the CIN high CNV LN clone (CNV clone C, blue), and CIN high CNV LVI clone (CNV clone B, pink), respectively. Once diverged, these clones appear to have undergone further copy number changes suggesting that while the CNV LVI clone and the CNV LN clone once shared a common ancestor, they evolved independently and gradually. By comparison, the CNV mixed-epithelial clone had a relatively flat evolutionary profile with only a few minor diverging subpopulations. These likely represent random changes in genetic copy number that ultimately were not advantageous to the tumor, nor did they promote further clonal outgrowth. A second maximum parsimony tree constructed based upon SNV identification showed a similar result whereby the SNV LVI clone (SNV clone B, pink) and the SNV LN clone (SNV clone C, blue) evolved separately and gradually from an ancestor of mixed-epithelial SNV clone origin (SNV clone A, green; Fig.4B). In contrast to the CNV phylogenetic tree, the evolutionary distance between the three SNV clones was found to be much smaller, and the SNV LVI clone appears to have arisen from a more closely related SNV LN clone. Altogether, this phylogenic analysis suggests that each of the CNV and SNV clones identified are evolutionarily distinct, and provides evidence for the LVI and LN clones being seeded by a mixed/epithelial-like clone originally present in the primary tumor of in this patient.

**Figure 4.**
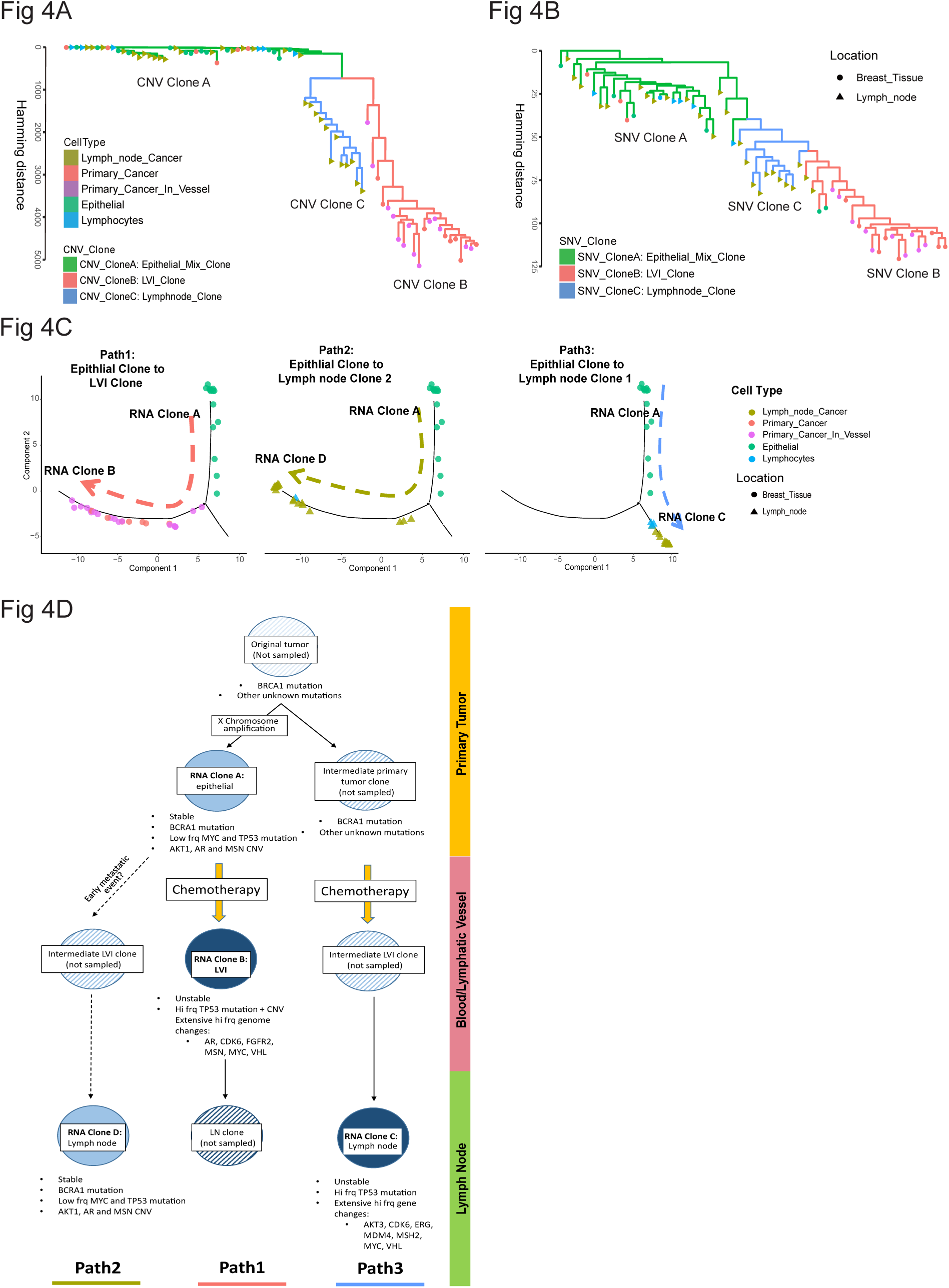
Evolutionary history of cancer cell clones reveals 3 distinct pathways of metastatic disease in one patient. Maximum Parsimony tree constructed based upon the CNV (A), and SNV (B), detection in cell clusters. Each marker represents an individual cell cluster isolated from either breast tissue () or lymph node tissue (). Markers are colored according to cell type. Lines between cell cluster markers correspond to CNV/SNV clone type and represent the evolutionary distance between cell clusters. (C) Metastatic transcriptional pathways of each RNA clone as defined by the Monocle 2.0 algorithm. Individual cell clusters are defined by cell type and tissue location as described in (A, B) and plotted against pseudotime. (D) Combined reconstruction of the evolutionary history of the tumor isolated from this patient. Cones detected by G&T-seq are depicted by solid circles while unsampled clones are represented with hash marks. Key genes potentially affected by oncogenic events are shown below each clone. Clone location is shown on the right (primary tumor, yellow; lymphatic vessel, pink; lymph node, green) and metastatic pathway (Path1, Path2, Path3) is depicted on the bottom.

Because chromosomal aberrations and genetic polymorphisms do not necessarily translate into predictable changes in gene expression, we assessed the evolution of our clones at a transcriptional level. In order to reconstruct the evolution of each RNA clone, we used the algorithm Monocle 2.0 to organize the cell clusters based upon their transcriptional similarity to build transcriptional trajectories representative of the gene expression pattern exhibited by individual cells clusters during their differentiation^35, 36^. Using this method, cell clusters were mapped against “pseudotime” which served as a quantitative measure for developmental progress. Branch points in these trajectories appeared when variations in gene expression were found to yield distinct differentiation events. Using the marker genes defined by the SC3 algorithm (Table S3.2), we plotted the RNA expression trajectories for each RNA clone using Monocle 2.0. This analysis revealed the presence of two transcriptionally distinct tumor cell differentiation fates or trajectories, each of which originated from cell clusters with an RNA epithelial clone phenotype (RNA clone A; Fig.4 C). Notably, the RNA LVI clone (RNA clone B, path1) and the RNA LN2 clone (RNA clone D, path 2) were found to differentiate along a similar trajectory (Fig.4C, Fig.S4), suggesting that they are related to each other developmentally. However, because the above tanglegram analysis found the RNA LVI clone to correspond to the CNV LVI clone exhibiting extensive chromosome aberrations, and RNA LN 2 clone mapped to the mixed-epithelial CNV clone with relatively few CNVs, it is not possible for RNA LVI clone and RNA LN2 clone to be genetically related. As such, it unlikely that the RNA LVI directly seeded the lymph node cancer clone as suggested by these trajectories. Thus, by combination analysis of DNA phylogenetic trees and RNA expression trajectories, we conclude there to be 3 clonal metastatic paths in this patient despite there being only 2 RNA trajectories. Pseudo-time gene expression profiles and Venn diagram analysis confirmed that these paths were largely unique with a distinct pattern of gene expression being associated with each path (Fig.S4, Table S3.2).

By combining these metastatic paths with oncogene assessment, we can begin to reconstruct an overall history of disease progression in this patient. As illustrated in Fig.4D, based on the high frequency of a *BRCA1* mutation, cancer cells in the original primary tumor likely had *BRCA1* driver mutation that was inherited by many of the existing cancer cells. Furthermore, because the X chromosome is amplified in Clone A (mixed-epithelial clone), clone B (LVI clone) and clone D (mixed-LN clone), but not Clone C (LN clone), Clone C, must have evolved from an ancestral clone present in the original primary tumor that either was not sampled, or is no longer present. Duplication of the X chromosome likely occurred later and led to the development the Clone A (mixed-epithelial clone) which is still present in the primary tumor, and metastasized to the patient’s lymph nodes as Clone D (mixed-LN clone). Based on the CNV phylogenic tree, Clone A (mixed-epithelial clone) also seeded Clone B (LVI clone) whose high mutational burden and chromosome instability are likely related to the chemotherapeutic regimen administered to this patient prior to tissue acquisition. These chemotherapeutic effects may have also played a role in the development of the genomically unstable Clone C (LN clone) from an unsampled primary tumor cell clone that lacked the X chromosome amplification and metastasized to the patient’s lymph nodes. In sum, with limited tissue sampling, G&T-seq allowed us to partially reconstruct a simplified model of tumor progression and metastasis in a TNBC patient with advanced, heterogeneous disease.

### Gene enrichment demonstrates distinct metastatic and immune related biological processes associated with each metastatic path

Having confirmed the existence of three transcriptionally distinct metastatic paths in this patient, we sought to determine the key biological processes and genes associated with each of these paths. Following gene-set enrichment of significantly changing genes associated with each pathway using Metascape^37^; Table S4.1), we selected the 20 most representative biological pathway groupings across all three metastatic paths and generated a heatmap to compare the relative significance of the selected biological processes to each metastatic pathway (Fig.5A). Similar to other reports, all three paths were enriched in the biological processes related to “epithelium morphogenesis”^38^, suggesting that genes associated with epithelial changes may have provided the foundation for metastasis in each of the clones identified. Furthermore, the metastatic pathways associated with genome instability (Path1, RNA LVI clone; Path 3, RNA LN1 clone), shared biological processes related to the cell cycle and oxidative stress, indicative of rapid proliferation and the involvement of hypoxia in the development of these clones. Metastatic pathways involving primary cancer cells (Path 1, RNA LVI clone; Path 2, RNA LN2 clone), were associated with biological processes related to “blood vessel development”, “epithelial cell migration”, and “regulation of cell adhesion”, all of which have been previously reported as related to tumor cell invasion and migration^39,40,41^. Additionally, each metastatic pathway was associated with unique biological processes such as “chromosome segregation” and “T helper cell differentiation” in path 1, and “extracellular matrix organization”, “toll-like receptor signaling” and “response to IFN-γ” in path 2. Path 3 by comparison was enriched with biological processes associated with immune regulation (“antigen processing and presentation, MHC I”), cellular metabolism (“oxidative phosphorylation”), and translation termination (Fig.5A, Fig.S5, Table S4.1).

**Figure 5.**
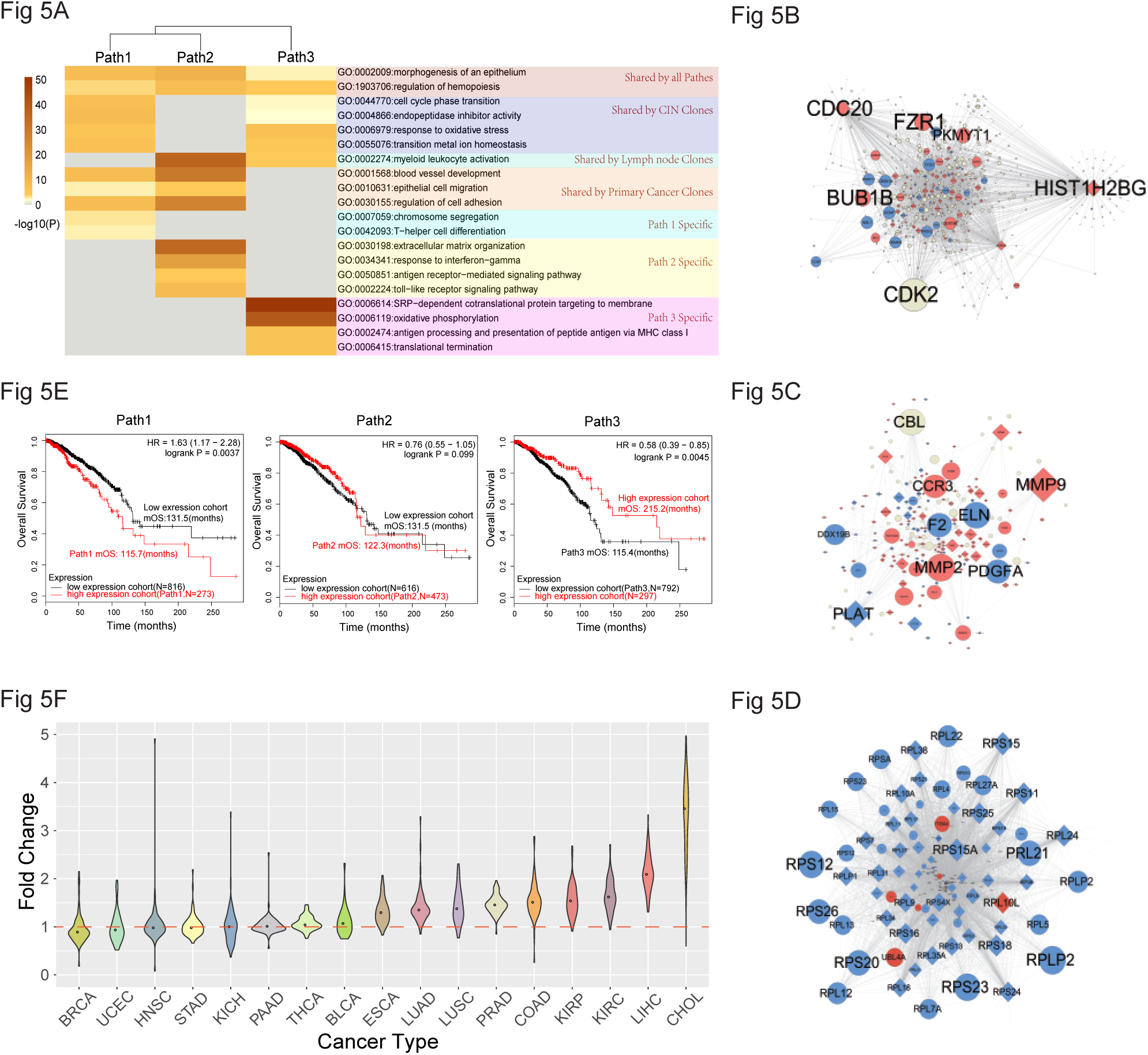
Biological process, hub gene, and overall survival analysis of each metastatic pathway reveals distinct gene signatures associated with variable patient outcomes. (A) Heatmap depicting the relative significance of the top 20 Metascape defined biological process groupings to each metastatic pathway. Biological processes are arrange based upon being: 1) shared by all paths, 2) shared by CIN clones, 3) shared by lymph node clones, 4) shared by primary cancer cell clones, or 5) path specific. Scale coloring represents the log P-value for the indicated biological process grouping. PPI network map of hub genes associated with path 1 (B), path 2 (C), and path 3 (D). Nodes represent individual genes, and edges denote a known association between genes. Red nodes represent genes exhibiting an increase in expression while blue nodes represent genes with decreased expression. Hub genes which were among top 10 hits from metastatic each path but not demonstrate a significant change in expression are depicted in grey. Node size denotes the relative importance of each gene to the network. (E) Kaplan-Meier plots depicting OS outcomes of breast cancer patients with high (red) vs low (black) expression for the top 20 hub genes associated with each metastatic path. OS curves were generated using KM-plotter from a cohort of 1089 breast cancer tumor samples and corresponding patient survival data. Hazard ratios and log-rank P-values are shown in the top right-hand corner of each Kaplan-Meier curve. mOS = median overall survival. (F) Violin plots depicting the fold change in expression of 80 RPL and RPS genes across 17 cancer types in comparison to normal tissue. RPL and RPS expression data was derived from the TCGA database incorporating ENCORI. Tumor types are sorted from lowest to highest median fold change in ribosomal protein expression. The median fold change value of each cancer type is presented by a dot in the middle of each violin plot. The horizontal orange dashed line marks a fold change value of 1.0. Violin shape corresponds to the density of data. A bigger bulbous in the violin means a higher density of the data and vice versa. The tumor types assessed are as follows: BRCA, Breast Invasive Carcinoma; UCEC, Uterine Corpus Endometrial Carcinoma; HNSC, Head and neck squamous cell carcinoma; STAD, Stomach adenocarcinoma; KICH, Kidney Chromophobe; PAAD, Pancreatic adenocarcinoma; THCA, Thyroid carcinoma; BLCA, Bladder Urothelial Carcinoma; ESCA, Esophageal carcinoma; LUAD, Lung adenocarcinoma; LUSC, Lung squamous cell carcinoma; PRAD, Prostate adenocarcinoma; COAD, Colon adenocarcinoma; KIRP, Kidney renal papillary cell carcinoma; KIRC, Kidney renal clear cell carcinoma; LIHC, Liver hepatocellular carcinoma; CHOL, Cholangiocarcinoma.

Having depicted the key biological processes of each metastasis path, we used the Cytoscape application CHAT ^42^, to facilitate the identification of “hub genes” (ie. highly interconnected genes critical to the biological system being studied) associated with each metastatic pathway, and to portray the corresponding protein-protein interaction (PPI) network. Hub genes associated with path 1 (RNA LVI clone) included those related to the cell cycle (*FZR*), cell division (*CDC20*) and nucleosome structure (*HISTIH2BG*; Fig.5B, Table S3.3). This is consistent with “Chromosome segregation” being identified as a key biological process associated with path 1, and suggests that chromosomal instability caused by abnormal chromosome segregation may have led to cancer metastasis by LVI associated clonal populations. Path 2 (RNA LN2 clone) was associated with inflammation and invasion related matrix metalloprotease (MMP) genes (*MMP9, MMP2*), as well as hub genes linked to cell adhesion *(VCAN*, *ITGB3*; Fig.5C; Table S3.3). The elevated expression of these genes, along with toll-like receptor and antigen receptor−mediated signaling being enriched biological processes (Fig.5A, Table S3.3), suggests that path 2 may be related the EMT/inflammation axis of cancer cell metastasis. Finally, an abundance of hub genes associated with path 3 (RNA LN1 clone) were found to be ribosomal proteins (48 of top 50 hub genes), all of which were downregulated with the exception of *RPL10L* (ribosomal protein L10 like; Fig.5D, Table S5.1). This reduction in the expression of transcripts encoding ribosomal proteins, combined with “antigen processing and presentation of exogenous peptide antigen via MHC class I, TAP-dependent” (GO: GO:0002479) and “translation termination” biological process, suggests that cancer cells associated with this pathway may be trying to limit protein synthesis and neo-antigen presentation on the cell surface.

### Pan-cancer analysis finds extensive down-regulation of transcripts encoding ribosomal proteins to be a potential marker of poor prognosis in breast cancer patients

As a final clonal assessment, we aimed to determine if the hub gene expression profiles associated with each metastatic path correlated to differences in breast cancer survival outcomes. To extrapolate our findings beyond this single TNBC patient, we employed the use of KM-plotter^43^ which utilizes the gene expression and survival data of cancer patients available from publically accessible databases to generate Kaplan-Meier overall survival (OS) curves. By comparing the survival outcomes of patients exhibiting high versus low mean expression of top 20 hub genes from each metastatic path, we found the hub gene expression profiles associated with path 1 (RNA LVI clone) and path 3 (RNA LN1 clone) to correlate with poor prognosis in breast cancer patients dichotomized by their respective low and high gene expression (Fig. 5E; *p* = 0.0037, HR 1.63 (1.17 – 2.28) and *p* = 0.0045, HR 0.58 (0.39-0.85), respectively). By contrast, differential expression of genes related to path 2 had little impact on overall breast cancer survival (*p* = 0.099, HR 0.76 (0.55 – 1.05)). Moreover, of the 1089 patient samples included in this OS analysis, a far larger number were found to exhibit a path 3 hub gene expression signature (N=792), in comparison to both path 1 (N=273) and path 2 (N=473; Fig. 5E). Given that increased ribosome biogenesis is a hallmark of many cancers and a recognized marker of poor disease prognosis^44^, our finding that extensive downregulation of both S ribosomal proteins (RPS) and L ribosomal proteins (RPL) associated with path 3 correlated with worse clinical outcomes was unexpected, and warranted further investigation. Of 80 ribosomal protein genes, 78 were found to be downregulated in path 3 (Table S5.1), and 64 of these had a log rank *P*-value lower than 0.05 when inputted into KM-plotter, correlating with poor prognosis in breast cancer patients when downregulated (Table S5.1). Only *RPL10L*, which exhibited increased expression in our path 3 hub gene analysis, was associated with improved prognosis (*p* = 3.2e-5).

To determine whether such broad ribosomal protein gene down-regulation occurs universally in breast cancer, or perhaps even in other cancer types, we performed a pan-cancer differential gene expression analysis of 80 RPL and RPS ribosomal protein genes across 17 different cancer types using The Cancer Genome Atlas (TCGA) expression data with integration from ENCORI (Encyclopedia of RNA Interactomes^45^; Table S5.2). Ribosomal protein genes were found to be broadly downregulated in TCGA breast cancer patients with 51 of 80 ribosomal proteins genes exhibiting significantly reduced expression in comparison to normal tissue (Table S5.3). Furthermore, breast cancer exhibited the lowest median fold change (<0.9) in ribosomal protein expression between tumor and normal tissues, when compared to 17 different cancer types (Fig.5F, Table S5.3). By contrast, 9 of 17 cancer types demonstrated extensive ribosomal protein gene upregulation with a median fold change > 1.1 (Table S5.3). Thus, not only does the hub gene expression pattern associated with metastatic path 1 (RNA LVI clone) and path 3 (RNA LN1 clone) correlate with a worse disease prognosis, reduced expression of ribosomal proteins could potentially serve as a marker of poor survival outcomes in a subset of breast cancer patients.

## DISCUSSION

Lymphovascular invasive tumor cells have received little attention despite their primary importance in determining the clinical outcomes of breast cancer patients. By combining the precision of laser capture microdissection with the sequencing power of G&T-seq, we were presented with a unique opportunity to use a multi-omic approach to explore the mechanisms promoting the metastatic spread of these invasive cells as they transited from the primary tumor to the axillary lymph nodes of a patient with TNBC. Our analysis found that LVI associated breast cancer cells exhibited common chromosomal and transcriptional features such as CIN. Moreover, we discovered evidence of polyclonal metastasis in this patient, with three transcriptionally distinct metastatic pathways identified. Genomic sequencing of the cancer cell clusters isolated further revealed a highly heterogeneous tumor harboring a vast mutational landscape, typified by extensive chromosome aberrations and single nucleotide mutations. These mutations affected numerous key oncogenes including *BRCA1, TP53, CHEK2, APC, MYC, CDK6* and *MCL1*, and were present not only in cancerous cells, but histologically normal breast epithelial cells as well.

While our ability to partially reconstruct the mutational and metastatic dynamics of such a complex disease stemmed primarily from the collaborative usage of G&T-seq and LCM, it’s important to note that the cell clusters analyzed in these experiments were dissected directly from H & E stained pathological slides prepared from frozen tissue samples. This approach enabled the histological identification and dissection of specific cells based on their association with specific morphological structures. Importantly, in harnessing this technique we were able to specifically capture clusters of lymphovascular invasive cancer cells directly from lymphatic vessels which, due to the unidirectionality of lymph flow, represent disseminated tumor cells categorically *en route* to the axillary lymph nodes from this patient’s primary tumor. Furthermore, because previous breast cancer studies have found LVI and lymph node metastasis to be mediated by collective cellular invasion^46^, the employment of LCM also allowed us to select and characterize more physiologically representative multi-cellular invasive units in our analysis.

By performing unsupervised hierarchal clustering of the cancer cell clusters isolated from this patient, we found LVI associated cells to be genomically and transcriptionally similar, exhibiting an exceptionally large number of chromosome aberrations, and expressing high levels of a number of genes related to invasion and EMT-regulation including *FOXC1, ART3, BIRC7, RAB40B, PTP4A3* and *NOTCH1*. This extensive CNV profile also suggests that the LVI associated cells isolated were genomically unstable, consistent with both *BCRA1* being a founding driver mutation in this patient, and the spindle checkpoint/chromosome maintenance genes being identified as key biological processes associated with this phenotype. Chromosome aberrations exist in a large proportion of human cancers^47^, and recent work by Bakhoum *et al*^48^ has linked CIN to metastatic spread whereby the rupture of CIN induced micronuclei elicits a cytosolic DNA response in breast cancer cells, ultimately converging on noncanonical NFκB activation concomitant with increased metastasis in animal models. While the number of NFκB and inflammatory genes directly associated with the LVI clones in our study is limited, we believe a similar mechanism may still be at play as RNA trajectory analysis placed LVI cells in state of differentiation preceding that of the lymph node cancer RNA clone D (RNA LN2 clone), which exhibited a clear inflammatory and invasive phenotype. Moreover, because the LVI associated cells in our study were isolated from within lymphatic vessels, their transcriptional profile may be suppressed as a means to circumvent immune surveillance and the physical stresses of lymphatic transit. It is also noteworthy to mention that a number of primary tumor cell clusters were found to exhibit a similar CNV/SNV profile as LVI associated cells, suggesting that these invasive cells may have arisen directly from cancer cells positioned along the stromal interface of the primary tumor. While spatial mapping of the cell clusters collected would be required to confirm this association, similar correlations have been described by other groups^49^, suggesting that these primary cancer cells could be therapeutically targeted to inhibit the metastatic spread of disease via the lymphatic system.

Our analysis also revealed evidence of polyclonal metastasis with two genetically and transcriptionally distinct cancer cell populations identified in the lymph nodes of this patient. Based on evolutionary evidence, these clonal populations were likely established by asynchronous metastatic events that occurred at both early (RNA LN2 clone) and late (RNA LN1 clone) time points in the tumor’s lifecycle. This is consistent in part with the theory that early dissemination can seed metastatic breast cancer^50, 51^. Moreover, these lymph node clones were associated with transcriptionally distinct pathways of metastasis, and disparate overall survival outcomes when extrapolated to a larger cohort of breast cancer patients with path 2 associated cells (RNA LN2 clone) exhibiting a relatively benign gene signature compared to cells from path 1 (RNA LVI clone) and path 3 (RNA LN clone 1). This is consistent with path 2 clones demonstrating a more pro-inflammatory phenotype that may encourage immune detection, but also points to a role for the EMT/inflammation axis in promoting disease progression and metastasis in this patient. Indeed, these possibilities are not mutually exclusive as immune surveillance mechanisms initially restricting tumor growth can be overcome by tumor induced immunosuppressive changes in the tumor microenvironment, and is well reviewed elsewhere^52,53,54,55,56^.

It is generally presumed that aggressive cancers have a high rate of protein synthesis. Accordingly, we were surprised to discover that Path 3 associated RNA LN1 clones displayed reduced expression of transcripts encoding numerous ribosomal proteins, and that this was associated with worse survival outcomes in breast cancer patients. Conversely, ribosomal protein expression is often increased in cancer and is generally regarded as a marker of poor disease prognosis^44^. However, coupled with “translation termination” and “antigen processing and presentation of exogenous peptide antigen via MHC class I, TAP-dependent” biological processes, we hypothesize that path 3 associated cells exhibit reduced ribosomal protein expression as a means of limiting aberrant protein production and neoantigen presentation on the cell surface. This could serve as both coping mechanism in response to proteostatic stress^44, 57^, and an immune evasion strategy to prevent cytotoxic T cell responses and immune cell infiltration of the tumor^58^. Moreover, this extensive downregulation in ribosomal protein expression and its effect on survival outcomes appears to be unique to breast cancer with a large proportion of other cancer types exhibiting increased expression of these genes. The mechanism behind this phenomenon, and its significance to breast cancer however, remains unclear. Because these experiments are a temporal assessment of an advanced heterogeneous disease, it is also uncertain whether this gene expression pattern is stable, or represents a temporary phenotype in the spectrum of tumor cell plasticity. Indeed, recent studies have shown that energetics and protein synthesis rates are highly plastic, and that alterations in these phenomena may underpin resistance to therapy^59^. Hence, further investigation into the role of ribosomal protein expression in breast cancer progression should be explored.

In summary, this research demonstrates the power of G&T-seq in assessing complex, heterogeneous diseases such as advanced metastatic TNBC when paired with LCM. With limited sample collection, we were able to reconstruct, to a great extent, the evolutionary history of a tumor isolated from a TNBC patient, which would not have been possible if single cell, genomic, or transcriptional analysis techniques had been employed alone. As exemplified by our results, cancer metastasis is truly a dynamic process and thus, no single use of whole genome, exome, or transcriptional sequencing techniques can be used to predict how a particular cancer cell will interact with its local microenvironment. As a consequence, multi-omic and anatomically defined assessments such as these may be required to accurately navigate the intricacies of tumor invasion and metastasis if a therapy for metastatic disease is to be realized.

## METHODS

### EXPERIMENTAL MODEL AND SUBJECT DETAILS

Tissue samples were provided by a 24 year old female with T3 N1 M1, ER(-), PR(-), HER-2(-), grade 3, invasive ductal breast carcinoma following palliative mastectomy and full axillary lymph node dissection. This patient had received previous treatment with two cycles of docetaxel and four cycles of doxorubicin/cyclophosphamide chemotherapy with little or no clinical response. At the time of sample collection, metastasis has occurred to the patient’s local regional axillary lymph nodes (3 of 15 nodes were positive for metastatic carcinoma), as well as the liver. Two tissue samples were collected from the tumor-stromal interface of the primary tumor, and two additional tissue samples were collected from carcinoma positive axillary lymph nodes. Samples were obtained within 10min from resection and immediately immersed in liquid nitrogen for flash freezing. The patient died ∼9 months following sample donation, and survived ∼18 months following initial diagnosis.

This study and analysis were approved by Health Research Ethics Board of Alberta, Cancer Care committee (Approval # HREBA.CC-18-0470), and the BGI institutional review board on bioethics and biosafety (Approval No. BGI-IRB 15148). Informed consent was obtained prior to the performance of experimental procedures.

### LCM and LCM cell clusters lysis

Frozen tissues were embedded in Clear Frozen Section compound (VWR cat no. 95057-838), sectioned on a cryostat (Leica, CM3050S) and stained with hematoxolin and eosin (H & E) to identify areas for laser capture microdissection (LCM). LCM was performed with a Leica CRT6000 Laser capture microdissection microscope (Concord, ON, Canada) within 15 min of sectioning to avoid RNA degradation. Cell clusters were classified as groups of cells over 10µm in diameter. Each cell cluster contained an estimated 50-200 cells. For LCM cell cluster lysis, extracted cell clusters were incubated with a cell lysis mixture (20µL RLT Plus buffer (Qiagen 1053393), plus 1µL of spike-in RNA (1:250,000)) and immediately placed on ice for subsequent steps.

### Oligo-dT30VN bead labelling and mRNA and gDNA separation

M-280 Streptavidin Dynabeads® (Invitrogen, catalog no. 11205D) were washed according to the manufactures recommendations, mixed 1:1 with biotinylated-Oligo-dT30VN primer (100μM), and incubated for 20min at room temperature. Oligo-dT30VN-labelled beads were then washed and resuspended in bead resuspension buffer (Superscript II first-strand buffer, RNase inhibitor, nuclease-free water (NF-H2O)). For mRNA and gDNA separation, 10μL of Oligo-dT30VN labelled beads was added to tubes containing lysed cell clusters. 2.5µL of RLT Plus buffer was added to each tube and the bead/cell suspensions were incubated for 20 min at room temperature. Samples were then placed in a magnetic for 1min for bead separation. The supernatants (containing gDNA), were transferred to a new tube. Oligo-dT30VN labelled beads were then washed twice with 10μL of G&T seq wash buffer (50 mM Tris-HCl, pH 8.3, 75 mM KCl, 3 mM MgCl2, 10 mM DTT, 0.5% Tween-20, 0.2× RNAse inhibitor), and the supernatants containing gDNA were transferred into new tubes and stored at −80°C until further processing. The remaining Oligo-dT30VN-labelled beads coupled to cell cluster extracted mRNA were prepared for reverse transcription.

### Reverse transcription, amplification, purification of cDNA

For reverse transcription (RT), 10µL of RT Mastermix (5×SuperScript II First-Strand Buffer, 5M Betaine, 100mM MgCl2, 100mM DTT, 100uM TSO, RNAse inhibitor, SSII, dNTPs (10mM)) was added to tubes containing the Oligo-dT30VN-labelled beads coupled to the mRNA from each cell cluster. Samples were placed in a Veriti™ 96-Well Thermal Cycler (Applied Biosystems, Catalog no. 4375786) for RT using the following program settings: 42°C 2min, 42°C 60min, 50°C 30min, and 60°C 10min. 12.5 µL of PCR Mastermix (2×KAPA HiFi HotStart ReadyMix, IS PCR Primer) was then added to each RT reaction tube. Tubes were placed in the thermal cycler for cDNA amplification at the following program settings: 98°C 3min, 98°C 20s, 67°C 15s, 72°C 6min, 98°C 20s for 22 cycles and 72°C 5min. Amplified cDNA was stored at −20°C until purification. To purify amplified cDNA, room temperature AMPure XP beads (Agencourt, Catalog no A63881) were added at a 0.8:1 ratio to cDNA containing tubes and incubated at room temperature for 5min. The supernatants were removed, beads were washed twice with 100µL 80% ethanol and allowed to dry. Beads were then resuspended in 21µL of NF-H2O.

### Tn5 cDNA library preparation and sequencing

1.0ng of cDNA from each sample was mixed with a fragmentation mixture containing the BGI enzyme Tn5 Transposase (BGI, catalog no. BGE005) embedded with adaptors, and heated to 55°C for 7 min. The reaction was stopped by adding 5 μL of 0.1% SDS to each sample. 25 μL of PCR reaction mix (5 x KAPA Fidelity Buffer, 10 mM each dNTP, PhoAd153 F primer (10 μM), Ad153 R primer (10 μM), Ad153-F-tag (0.5 μM), Ad153-R-tag (0.5 μM), KAPA HiFi DNA polymerase), was added to each fragmented cDNA sample. Samples were transferred to a thermal cycler for amplification using the following program settings: 72°C 5 min, 95°C 3 min, 98°C 20s, 60°C 15s, 72°C 25s for 15 cycles, 72°C 5min. After amplification, 0.6X and 0.2X AMPure XP beads were used to select 300bp±100bp size fragments. These were then pooled for a total of 520ng cDNA per sample. cDNA in was cyclized by adding 20 µM Splint oligo (Invitrogen, Shanghia, China), and NF-H2O to each sample for a final volume of 70μL and heating samples to 95°C for 3min. 10x TA buffer (100mM ATP, T4 DNA ligase (600U/µL), NF-H2O), was added to each reaction for a final volume of 120μL. Samples were incubated at 37°C for 1hr. EXO digestion was then performed by adding 10x TA buffer mixed with EXO I (20U/µL), EXO III (100U/µL) and NF-H2O to each sample for a final volume of 128μL. Samples were incubated at 37°C for 30 min. Reaction products were then purified by adding 320μL of AMPure XP beads to obtain the cDNA library. Rolling circle amplification (RCA) was performed to produce DNA Nanoballs (DNBs) that were loaded on to the BGISEQ-500 sequencing platform (BGI, Shenzhen, China). Qualified cDNA libraries were sequencing with 100bp paired-end reads.

### Purification and amplification of gDNA

To enriched cell cluster gDNA, AmPure XP Beads were added to gDNA containing supernatants at a 0.6:1 gDNA to bead ratio, and incubated for 8 min at room temperature. The supernatants were then discarded and the remaining beads were washed twice with 100µL 80% ethanol, followed by the addition of 5µL NF-H2O. gDNA was amplified using a REPLI-g Single Cell Kit (Qiagen, Catalog no. 150345). Briefly, 3.5µL of Buffer D2 (denaturation buffer) was added to the beads and incubated for 10 min at 65°C. The reaction was stopped by adding 3 µL of Stop Solution to each sample. 40µL of Master Mix (29µL REPLI-g sc Reaction Buffer, 2µL REPLI-g sc DNA Polymerase, NF-H2O sc), was added to each denatured DNA sample for amplification. Samples were then incubated at 30°C for 8 h and the reaction was stopped by heating samples to 65°C for 3 min.

### Housekeeping test of MDA products

Prior to the WES/WGS sequencing, the quality of the amplified DNA products was assessed using a multiplex PCR based method that evaluated the presence of 8 genes (CYB5A, PRPH, GABARAPL2, ACTG1, NDUFA7, UQCRC1, MYC, MIF) from different chromosomes. 1µL of PCR mix (3.0 µL 10x Buffer, dNTP (2.5mM) 3.2 µL, 3.0 µL Primer Mix (10μM) 0.2 µL 100 x BSA, 0.4 µL rTaq (5U/µL)), was added to each amplified gDNA sample. These were then placed in a thermal cycler for amplification of the above gene products using the program settings: 95°C 4min; 95°C 30s; 56°C 50s and 72°C 1min for 35 cycles; 72°C 10min. Agarose gel electrophoresis was then performed on the PCR amplification products. Samples in which 4 or more bands were detected were subjected to downstream library preparation.

### WGS library preparation and sequencing

Whole genome sequencing (WGS) libraries were constructed from quantified and amplified gDNA from each cell cluster using an MGIEasy DNA Rapid Library Prep Kit (BGI, catalog no, 940-200033-00), and the BGISEQ-500 sequencing platform. Briefly, high-quality gDNA was randomly fragmented using a Covaris LE220 ultrasonicator (Covaris, Woburn, MA, USA). AMPure XP magnetic bead-based cleanup was conducted to select fragments ranging from 100-700 base pairs (main band 200-300bp). Selected fragments were tailing end repaired by adding Adapter Mix after which the ligated was DNA purified. Purified DNA samples were transferred to a thermal cycler and amplified with the following program settings: 95°C 3 min; 8 cycles of 98°C 20 sec, 60°C 15 sec, 72°C 30 sec; then 72°C 10 min. The PCR products were purified with AMPure XP magnetic beads. Samples were mixed with different barcodes and NF-H2O was added to each sample for a final volume of 48μL. The homogenized PCR products were then denatured by heating them to 95°C for 3 min in a thermal cycler. 11.8 μL of Reaction Mixture was added to each sample. Denatured DNA was circularized by placing samples in a thermal cycler for 30 min at 37°C. Circularized DNA was then digested by added 4µL of Digestion Reaction Solution to each sample for 30 min at 37°C. This reaction was stopped by added 7.5 μL of Digestion Stop Buffer. Single stranded circular DNA was then purification using AMPure XP magnetic beads. RCA was performed to produce DNBs that were loaded on to the BGISEQ-500 sequencing platform. The qualified WGS libraries were sequenced with an average coverage of 0.5∼1X with 100bp single-end reads.

### WES library preparation and sequencing

Whole exome sequencing (WES) libraries were constructed from quantified and amplified cDNA from each cell clusters using MGIEasy Exome Capture V4 Probe set (BGI, Shenzhen, China). cDNA pre-hybridization was performed by heating samples to 95°C for 5min followed by hybridization at 65°C for 24h. After elution of hybridized cDNA products, a post-PCR reaction mixture (2X KAPA HiFi HotStart Ready Mix, Ad-153-F (20μM) and 4 NF-H2O) was added to each sample. Samples were divided in half and amplified on a thermal cycler with the following program settings: 95°C 3 min;13 cycles of 98°C 20 sec, 60°C 15 sec, 72°C 15sec; then 72°C 10 min. PCR products were then purified using AMPure XP magnetic beads. PCR products totaling 330ng were pooled together. Samples were processed for splint circulation and made into a single strand circular DNA for WES library construction. RCA was performed to produce DNBs that were loaded onto the BGISEQ-500 sequencing platform and sequenced for 1 lane with 100bp paired-end reads.

### WGS data processing and CNV calling of cell clusters

Deconvoluted sequencing FASTQ data corresponding to each cell cluster sample was aligned to HG19/NCBI37 using BWA-MEM algorithms (BWA, Version: 0.7.17). SAMtools (Version: 0.1.19) was used to sort BAM files, mark and removed PCR duplicates, and calculate each chromosome’s depth and coverage. BAM files produced by alignment were counted in 5k,10k, 20k, 50k genomic bins using a “non-overlapping” “variable binning” strategy as previously describe^25, 60^. ‘“Variable binning” results in each bin having variable start and end coordinates. Since the reference length is fixed, a lower genome cutting bin number will lead to a border bin length, with the median genomic length spanned by each bin for 5k, 10k, 20k, 50k cutting bins being 554kb, 220kb, 136kb and 54kb, respectively. The variable start and end coordinates were determined by mapping back 200 million simulated sequence reads with 100nt length to the HG19/NCBI37 reference to determine bins for further calculation. ‘Non-overlapping bins’ means the boundary of each bin did not overlap with the genome coordinates, enabling us to clearly identity the copy number variation value of each chromosome segments and annotate the CNV affected genes. Unique normalized read counts of each variable bin was calculated using the Circular Binary Segmentation (CBS) method from R Bioconductor ‘DNAcopy’ package ^61^. The parameters used for CBS segmentation were alpha = 0.05, nperm=1000, undo.SD=1.0, min.width=5. Default parameters were used for MergeLevels which removed erroneous chromosome breakpoints. The median absolute pairwise difference (MAPD) was calculated to quantify the copy number noise of each cell cluster. We choose the 5k bin for further clustering analysis as it had a higher average MAPD value compared with 10k, 20k and 50k bins. Next, we filtered out cell clusters with coverage lower than 10% as calculated using BEDTools (v2.17.0), and with a MAPD greater than 1.00. This accounted for approximately 18% (17 of 97 cell clusters) of the total cell clusters for this patient.

### CNV heatmap construction and clone identification

To construct a clustered CNV heatmap, we calculated Euclidean distances from the copy number data matrix. Each column represents one cell cluster and each row represents the relative copy number ratio of diploid cells from each segment. Ward. D2 hierarchical clustering algorithm was performed in R using the pheatmap (version: 1.0.12) package available on CRAN. Columns representing a single cell were hierarchically clustered using Ward.D2 linkage on the basis of pairwise Euclidean distances, and the x-axis was ordered by chromosome coordinates of genome position. To estimate the reliability of each clade defined by the Ward.D2 clustering algorithm, we used the bootstrap method by R package pvclust (Version: 2.0.0) to calculate the AU (Approximately Unbiased) p-value and BP (Bootstrap Probability) value. An AU p-value > 0.95 is seen as stable clade to define a CNV clone.

### Maximum Parsimony tree construction

To detect common chromosome breakpoints and segments that were shared by cell cluster samples in each identified DNA-CNV clone, we applied a multiple-sample population segmentation algorithm using a Bioconductor R package copynumber (Version: 1.22.0), with parameter gamma=1. Piecewise constant curves were fitted to the cell cluster CNV data by minimizing the distance between the curve and the observed multi-cell data, and returning multi-cell segments with a fitting clonal CNV result. The Maximum Parsimony tree was calculated from the CNV-clone matrix using the parsimony ratchet algorithm with R package phangorn (Version: 2.4.0). Homozygous deletion, heterozygous deletion, neutral, or amplification, were treated as characters, and missing values were treated as ambiguous items. Hamming distance was calculated for branch lengths with R package ape (Version: 5.1). Phylogenetic trees were exported in Newick format, and R package ggtree (Version: 1.14.6) was introduced for visualization.

### WES data processing, SNV clone calling, heatmap construction, and clone identification

On target sequencing reads in the FASTQ files of each cell cluster were cleaned by SOAPnuke (Version: 1.5.6), aligned by BWA (Version: 0.7.17), sorted by SAMTools (Version: 0.1.19), PCR duplications removed by Picard, and realigned by GATK Realigner. The putative SNVs for each cell cluster were called by Monovar with the parameter setting: p=0.002, a=0.2, t= 0.05, m=4, c=1 and annotated by ANNOVAR (humandb:20170901). Putative SNVs were then filtered using the following parameters: 1000G_ALL<0.5%, ExAC_ALL<0.5%, ESP6500siv2_ALL<0.5%, genomicSuperDups score<0.9. Only mutations with nonsynonymous, stop/gain and stop/loss in exonic and splicing regions were kept as final SNVs for each cell cluster. The SNV matrix containing the final SNVs of each cell cluster was construed, where 0 represent not mutated or unidentified SNV sites, 1 represent heterozygous SNV sites and 2 represent homozygous SNV sites. SNV heatmaps were constructed using the Ward.D2 clustering method with Euclidean distances of SNV matrix construed by Monovar. DNA-SNV clones were identified using the same method as described above for the DNA-CNV clones.

### WTS data processing and quality control

Sequencing data was first processed to filter out low quality reads which were defined as: 1) ‘‘N’’ bases accounting for 5% read length; 2) Bases with quality < 15 accounting for 50% read length; 3) Containing the adaptor sequence; 4) Duplicate reads. The reads that passed were then aligned to ribosomal RNA sequences downloaded from NCBI Reference Sequence Database using SOAPaligner (soap2 V2.21t). The unmapped reads were aligned to human genome assembly GRCh37 (hg19). Gene expression TPM was calculated using bowtie2 plus RSEM with default parameters. Saturation curves were then calculated for each cell cluster, and curve densities were compared between each saturation curve. Cell clusters showing an unsaturated curve and an obviously skewed density plot were considered unqualified samples.

### RNA clone calling of cell clusters

For each cell cluster, TPM was calculated for each given gene in Refseq. TPM matrices for genes were supplied to SC3^30^, a single-cell consensus clustering pipeline, with the following parameters: pct_dropout_min=2 and pct_dropout_max=90. After the consensus matrix was built by SC3, the average silhouette width and stability index values were calculated. These were combined with cell type and the best empirically performing clustering were determined. Once the stable clusters were determined to identify RNA clones, genes exhibiting the highest variability amongst each LCM cell cluster was calculated. The resultant marker genes were identified by a ROC curve (AUROC) > 0.85 and p-value < 0.01. The top ten maker genes of each cluster were shown in the heatmap with a log2(TPM+1) value.

### RNA trajectory reconstruction and gene set enrichment

TPM marker genes identified using SC3 we supplied to Monocle2 to generate pseudotime plots which reflect cell fate decisions and differentiation trajectories. Genes were identified as being differentially expressed between trajectories using a cut-off q value of q<0.01. We choose the top 800 qualifying genes and defined these as “significant changing genes” for each RNA trajectory. By further analyzing the branches of each RNA trajectory, we found statically significant (q<0.001) branch-dependent genes. We used the previously defined “significant changing genes” and the branch-dependent genes associated with each trajectory to do GSEA, and to determine the related GO BPs (gene ontology biological processes). We defined significant biological processes as those with a q<0.01 as calculated by GSEA.

### DNA and RNA-clone comparison

The cell cluster DNA-CNV-tree was constructed using the Euclidean distance of the CNV data matrix, clustered with the hclust function using WARD.D2 linkage in R, then outputted as Newick format. Each clade of the tree reflects a DNA-CNV-clone. The RNA-tree was constructed by SC3 and the tree clade was outputted as Newick format. We mapped the DNA-CNV-tree and DNA-SNV-tree using Tanglegrams generated by the dendextend (version: 1.8.0) package in R. Samples found on only one side of the tree were remove from our analysis. The DNA-CNV-tree was mapped to the RNA-tree to compare the consistency between DNA and RNA clones by the same process.

### Gene Set Enrichment and Hub Gene Identification

Genes exhibiting a significant change in expression (Table S3.2,Top 800 with q<0.01), from each metastasis paths identified following Monocle 2.0 analysis was supplied to Metascape ^37^ to independently perform biological pathway and process enrichment analysis using the following ontology sources: GO Biological Processes and GO Molecular Functions (Database Last Update Date: 2019-06-11). All genes associated with each metastatic path were used for this enrichment. Enrichment terms with a p-value < 0.01, a minimum count of 3, and an enrichment factor > 1.5, were collected and grouped into clusters based on membership similarities calculated by Metascape. More specifically, accumulative hypergeometric distribution and Banjamini-Hochberg were applied to calculate the p-values and q-values of each term^62^. Kappa scores were used as a similarity metric when performing hierarchical clustering on the enriched terms, and sub-trees with a similarity of > 0.3 are considered a cluster. The most statistically significant terms within a cluster is chosen to represent the each cluster. To further capture the relationships between terms, a subset of enriched terms was selected and used to generate a network map where by terms with a similarity > 0.3 are connected by edges. Networks were visualized using Cytoscape with a layout generated by employing the yFiles Radial method, where by each node represents an enriched term and is colored by its cluster ID. We further grouped each term into five subgroups (“shared by three paths”,” Shared by Primary Cancer Clones”, “Shared by CIN Clones”, “Shared by Lymph node Clones” and “Path Specific”), and marked these subgroupings on each Cytoscape network based upon cell cluster location. We then choose the 20 most respective terms combined across all metastatic paths to generate a heatmap in which the coloring is representative of the log p-values associated with each term for each metastatic path. Terms in the heatmap were further defined as belonging to 1 of 3 three categories: 1) Metastasis related terms”; 2) “Immune related terms”; and 3) “Other terms”.

### Hub gene identification and Kaplan-Meier analysis

Genes exhibiting a significant change in expression, as well as their fold-change value (Table S7), were supplied to the Cytoscape application CHAT (Contextual hub analysis tool, version:1.0.5) to identify hub genes related to each metastatic path, and generate a PPI network from the BioGRID database of Human/Homo sapiens Taxonomy. The mean expression of the top 20 hub genes for each metastatic path were also supplied to Kaplan-Meier Plotter^43^ to generated Kaplan-Meier survival curves for these signatures, based upon the survival data of 1089 breast cancer patients sourced from GEO (Gene Expression Omnibus), EGA (The European Genome-phenome Archive), and TCGA (The Cancer Genome Atlas) databases. The top 20 genes were selected based upon the significance of prognosis prediction. 80 of the most characterized ribosome protein genes (RPS and RPL) were also supplied to Kaplan-Meier Plotter to generate Kaplan-Meier overall survival p-value across 21 types of cancer available in Kaplan-Meier Plotter database (TableS5.4).

### Pan-cancer ribosome protein gene analysis

80 of the most characterized ribosome protein genes (RPS and RPL) were individually supplied to ENCORI (The Encyclopedia of RNA Interactomes; http://starbase.sysu.edu.cn/index.php), a Pan-Cancer Analysis Platform which enables differential gene expression analysis between tumor and normal tissues using available mRNA data from TCGA (The Cancer Genome Atlas; https://www.cancer.gov/about-nci/organization/ccg/research/structural-genomics/tcga). The differential expression data of these 80 ribosomal proteins was available in ENCORI for 17 types of cancers with 7086 tumor and 704 normal samples. Results of this analysis are shown in TableS5.2.

### DNA and RNA Amplification and Library Qualification

Amplification DNA was quantified using Housekeeping genes as described using a Qubit dsDNA High Sensitivity kit(Invitrogen, USA; catalog number: Q32854) as per the manufacturer’s instructions. Amplified RNA was quantified using a Agilent’s 2100 Bioanalyzer (Agilent Technologies, CITY, STATE) and the library was quantified using Qubit dsDNA High Sensitivity kit (Invitrogen，USA；catalog number: Q32854).

### Statistical analysis

The statistic of AU (Approximately Unbiased) p-value is calculated by multiscale bootstrap resampling by pvcluster package (v2.0.0). P-values of chi-square test were based on asymptotic theory. Silhouette width and stability index statistics were calculated using the SC3 package. All p-values were two-sided and q < 0.01 was considered significant. Maker genes of each RNA clonal were defined by a q-value <0.01 and a ROC value >0.85. A Significant change in gene expression was defined as q<0.01 with top800 genes of three paths. The Hub gene for each paths were defined using an adjust P-value<0.01. All other statistical analyses were carried out as described in the text using the R statistical environment (v3.4.4 and v3.5.0). Pathway network graphs were generated using Cytoscape (v3.6.1).

## DATA ACCESS

All raw and processed sequencing data generated in this study are available in the CNGB Nucleotide Sequence Archive (CNSA: https://db.cngb.org/cnsa; accession number CNP0000440).

## ACKNOWLEDGEMENTS

This research was supported by the Science, Technology and Innovation Commission of Shenzhen Municipality under grant No.GJHZ20170314152701465 and Joint Fund of the National Natural Science Foundation of China and Natural Science Foundation of Guangdong Province (U1601224). This research was initiated with funding by Alberta Innovates, in the form of an AITF/iCORE Strategic Chair (RES0010334) to G.K.S.W. We would also like to thank Dr. Ivan Topisirovic for his input and comments on the manuscript.

## AUTHOR CONTRIBUTIONS

G.K.S.W., J.M., B.L., Y.H., and X.Z. conceived the study. G.K.S.W., and J.M. designed the experiments. A.J., G.L., S.S. collected clinical samples. W.W., L.F., Q.Z and T.J. performed the experiments. Z.Z., Q.S., C.C., X.Z analyzed the sequencing data. D.L.H., G.K.S.W., Z.Z., J.R.M., L.M.P., wrote the manuscript with all authors contributing to writing and providing feedback.

