## Supplemental Figures for "Genome profiles of lymphovascular breast cancer cells reveal multiple clonally differentiated outcomes with multi-regional LCM and G&T-seq" (from another version of this preprint: version 1, 2019-10-17)

**Supplemental Figures outline:**

Figure S1. Cell cluster LCM figures and corresponding DNA CNV profile.
Figure S2. SNV heatmap of 1744 non-filtered SNVs, and comparison of heterozygous and homozygous mutation burden between the three SNV clones.
Figure S3. Venn diagrams comparing the number of genes affect by CNV, SNV, and RNA expression changes.
Figure S4. Venn plot analysis of RNA clones, pseudo-time trajectory, and corresponding gene heatmap trajectory for the three metastasis paths.
Figure S5. Enriched biological process networks for the three metastasis paths.

Figure S1 Cell cluster LCM figures and corresponding DNA CNV profile

Note: Microscopy figures of H&E stained pathological frozen tissue sections before and after the removal of cancer cell cluster using LCM. The corresponding copy number variation profile for each cell clusters is also presented. The 80 LCM cell clusters which passed WGS QC are depicted.

Figure S1.1 LCM clusters from CNV Clone A epithelial.

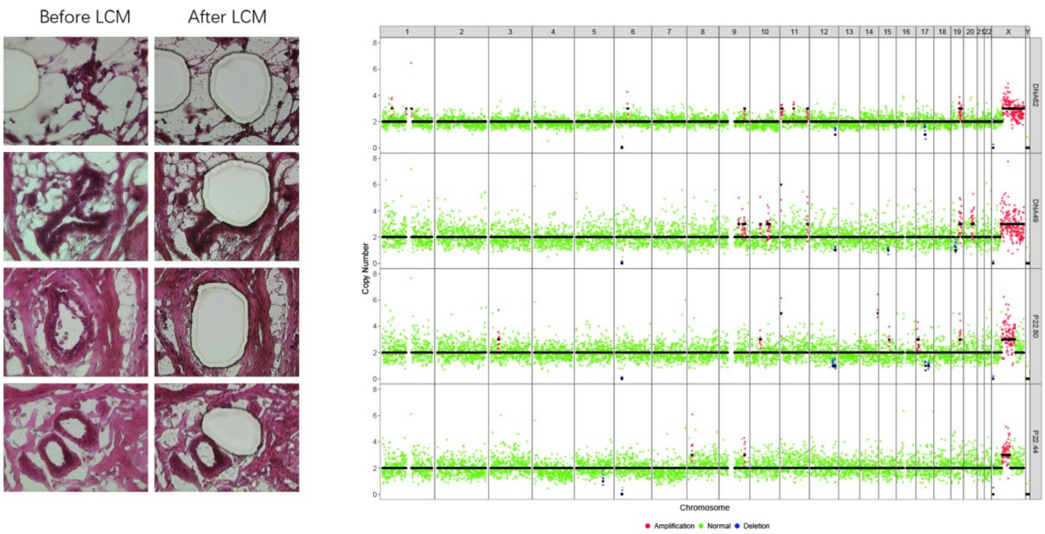


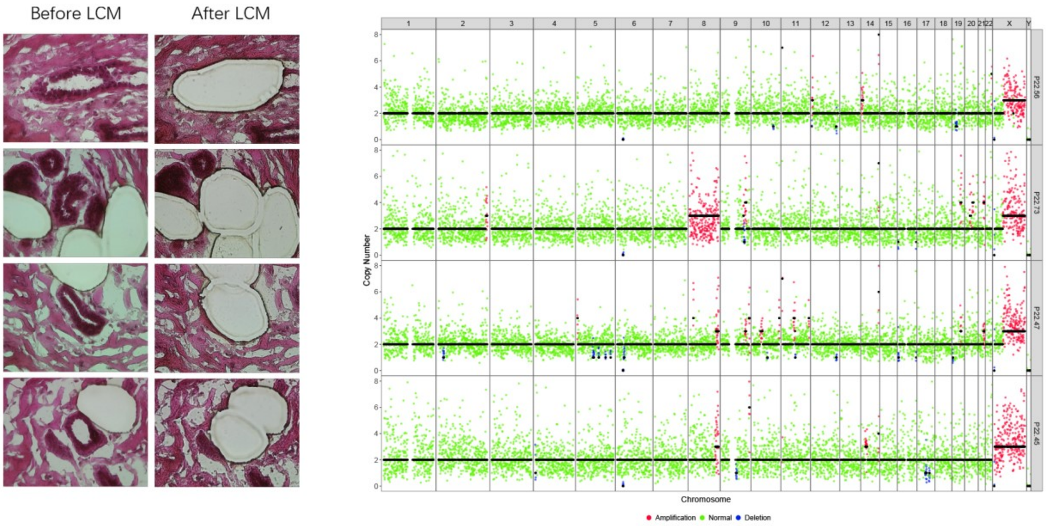


Copy Number Abbreviations for Clone A Epithelial

Figure S1.2 LCM clusters from CNV Clone A epithelial.

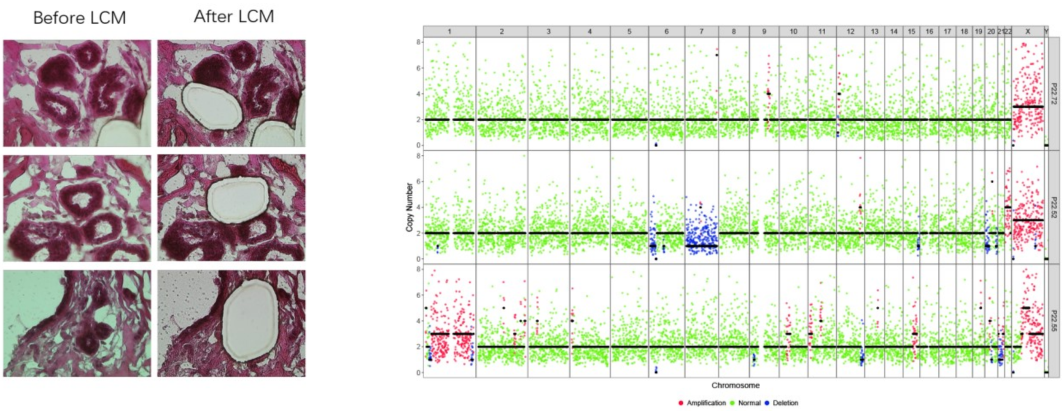


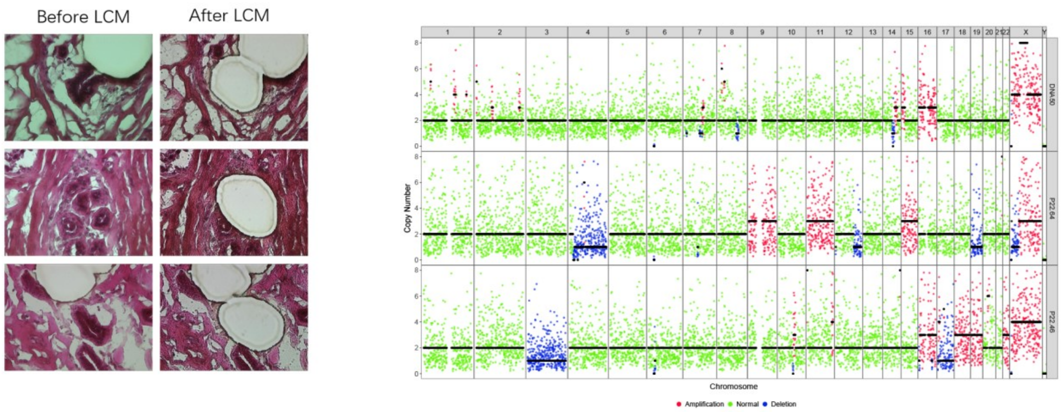


Copy Number Abbreviations for Clone A Epithelial

Figure S1.3 LCM clusters from CNV Clone A lymphocytes

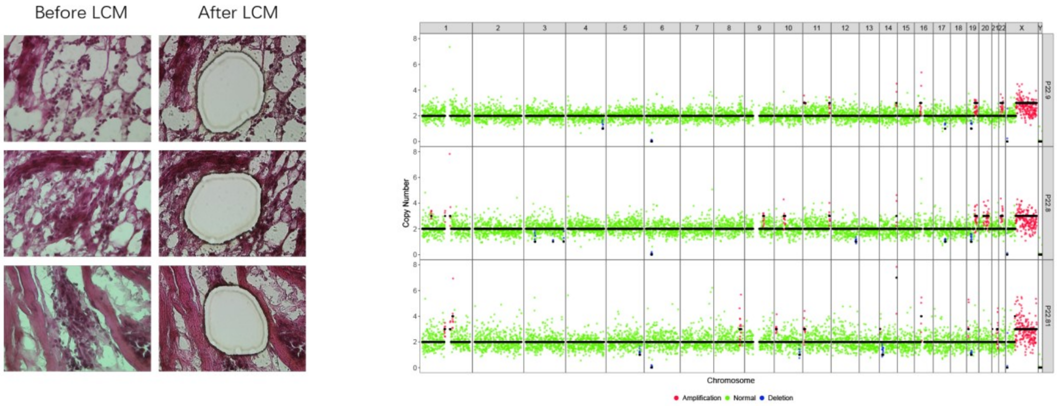


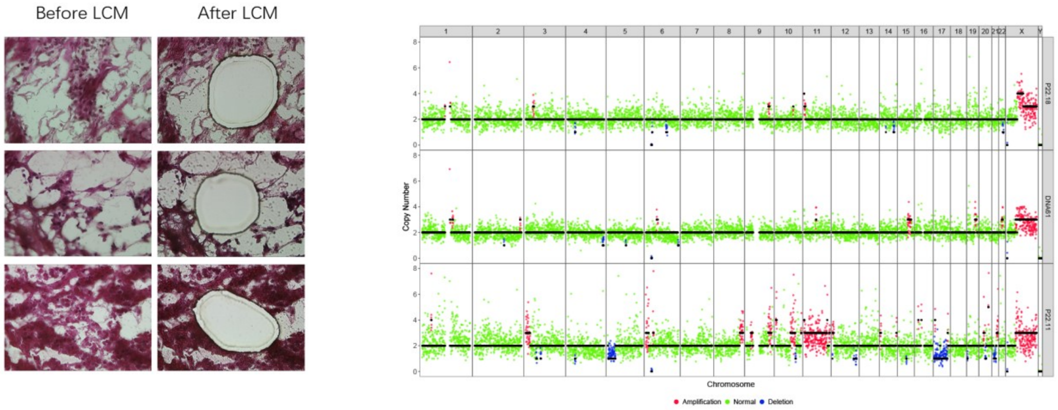


Copy Number Abbreviations for Clone A Lymphocytes

Figure S1.4 LCM clusters from CNV Clone A primary cancer.

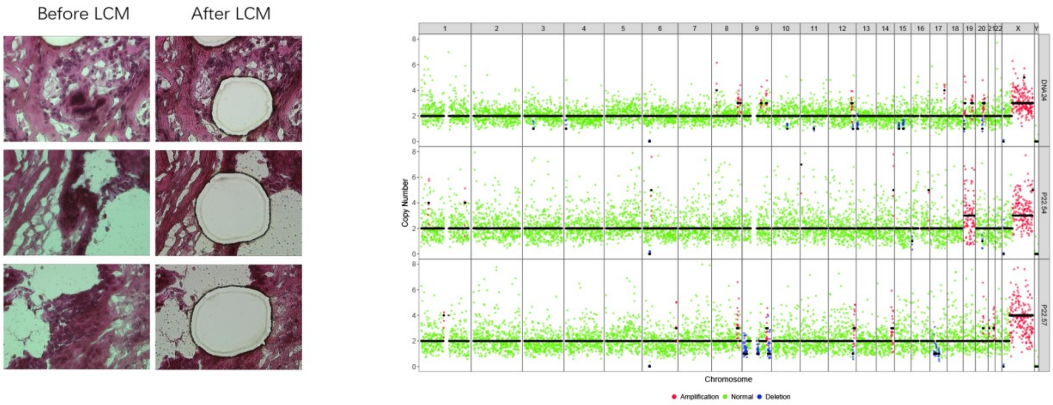


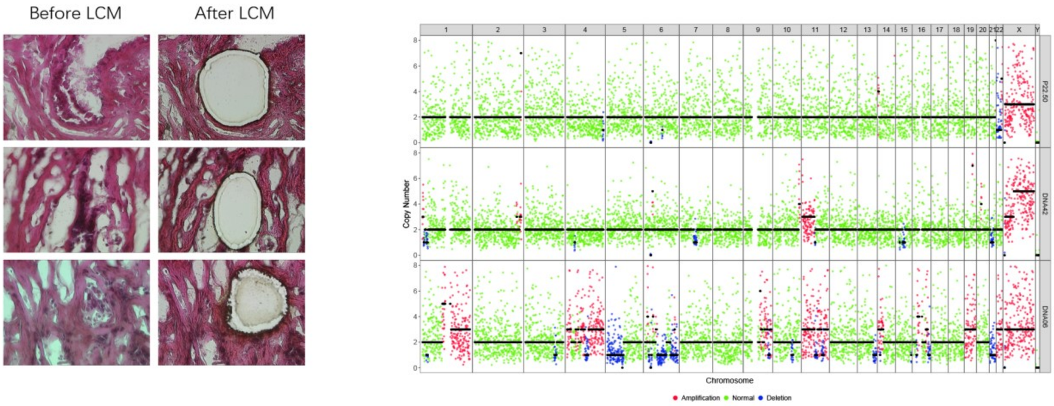


Copy Number Abbreviations for Clone A Primary Cancer

Figure S1.5 LCM clusters from CNV Clone A lymph node cancer

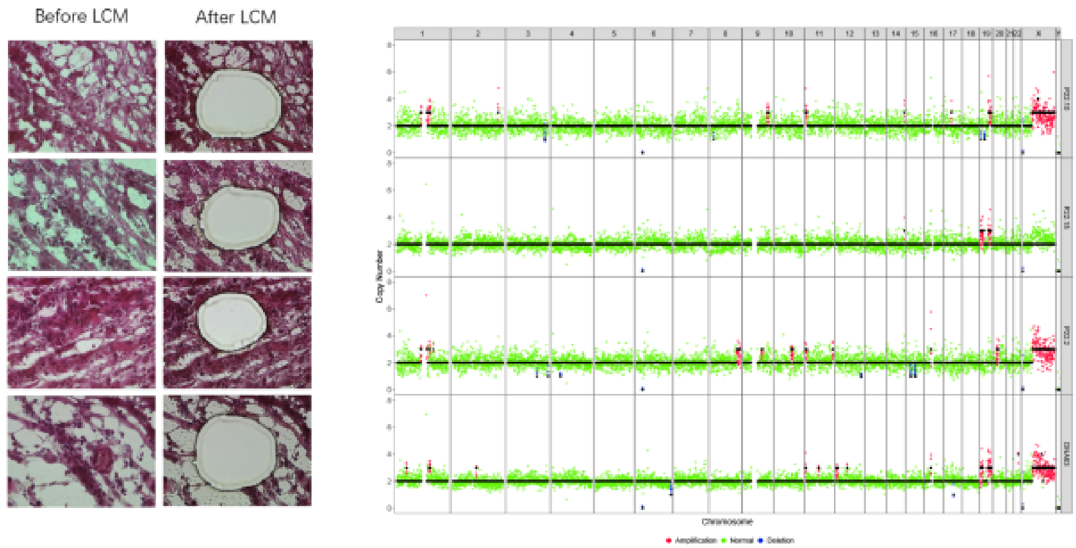


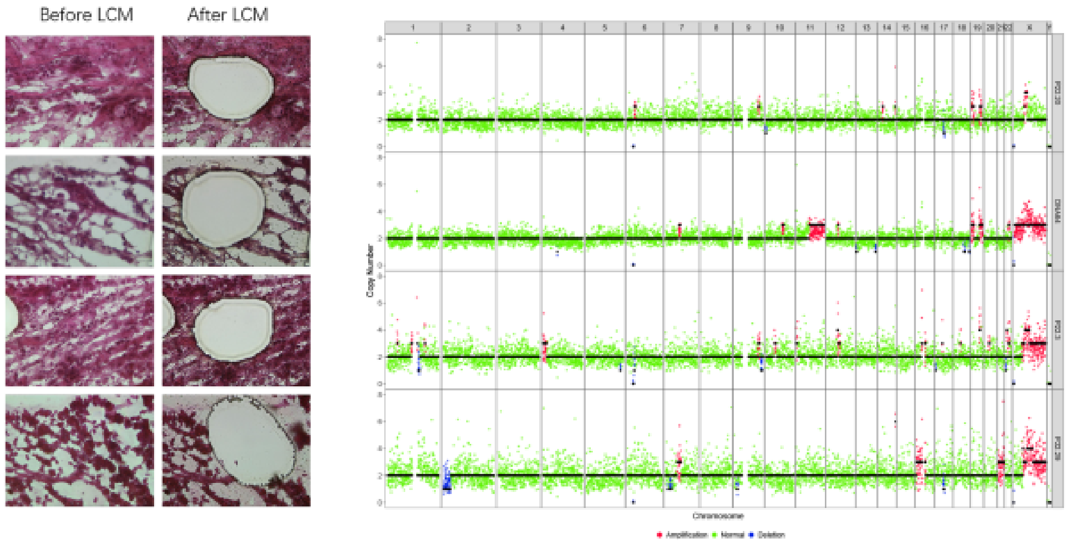


Copy Number Abbreviations for Clone A lymph node Cancer

Figure S1.6 LCM clusters from CNV Clone A lymph node cancer

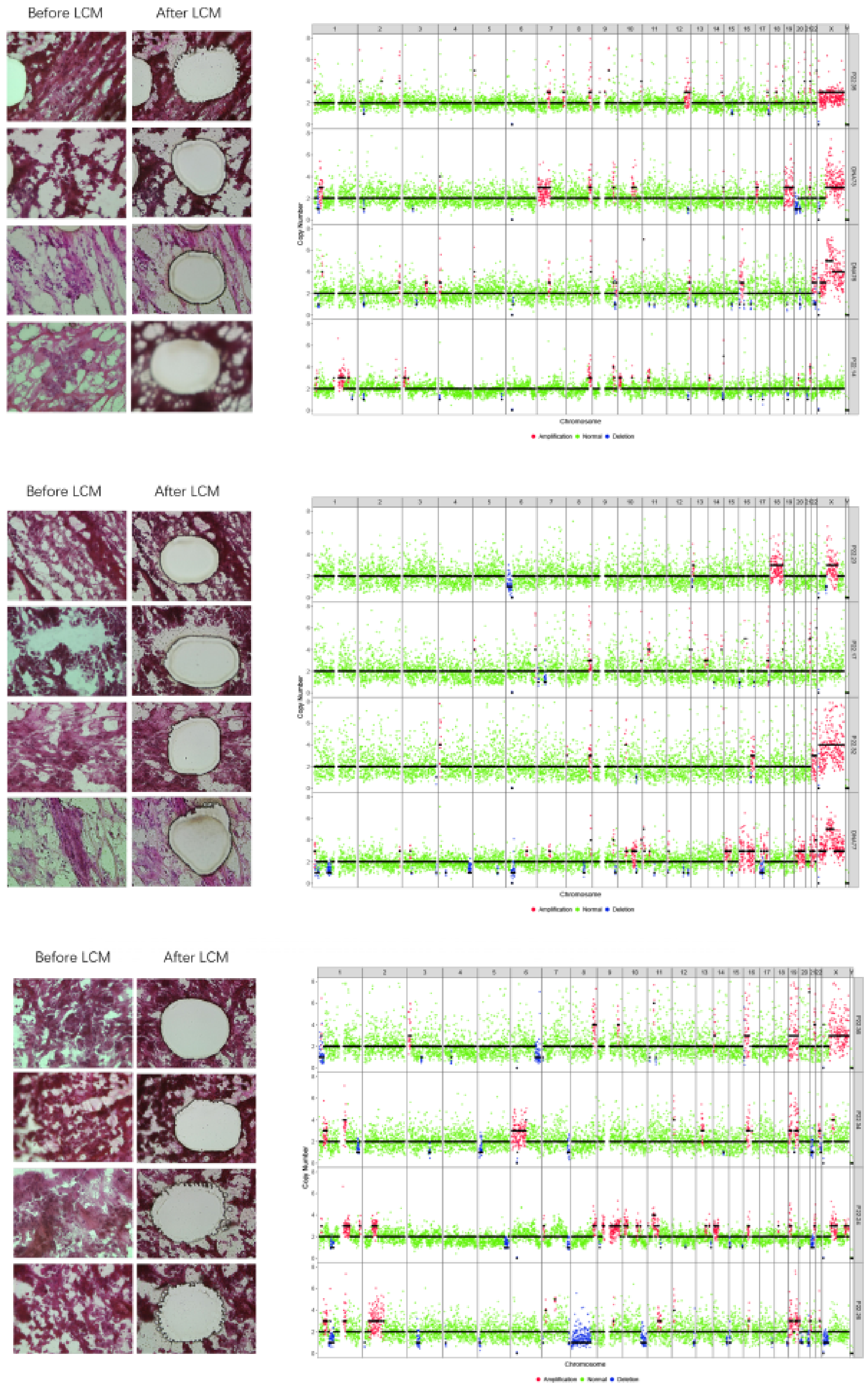


Copy Number Abbreviations for Clone A lymph node Cancer

Figure S1.7 LCM clusters from CNV Clone B primary cancer

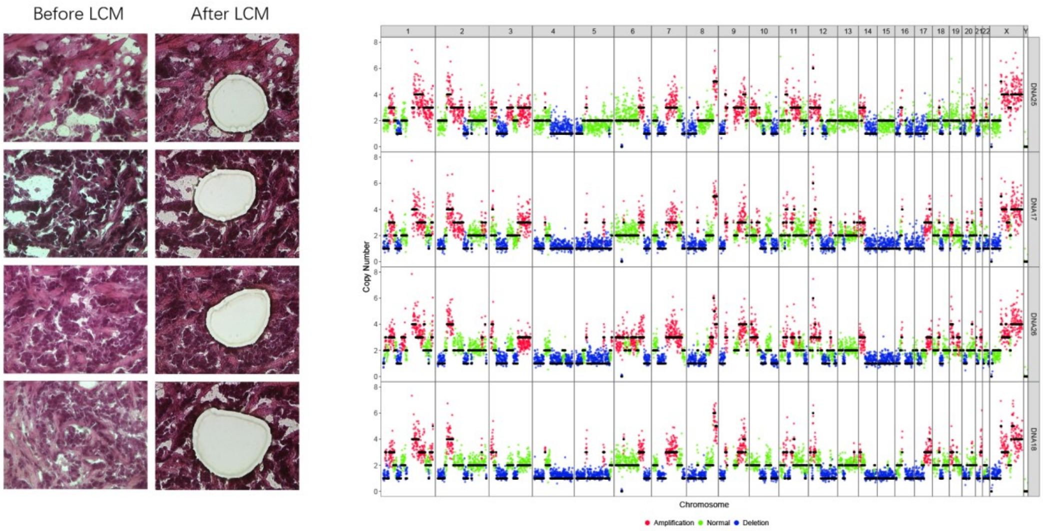


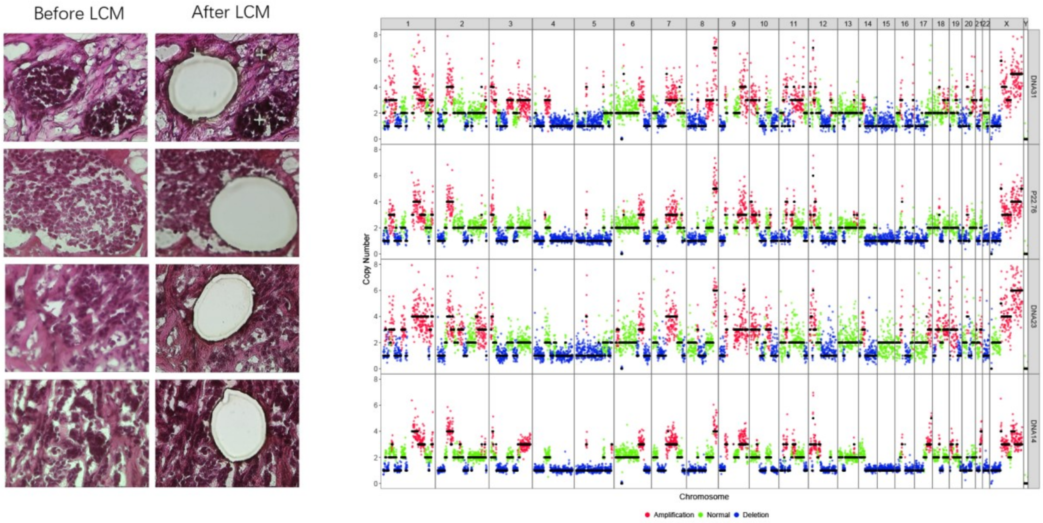


Copy Number Abbreviations for Clone B Primary Cancer

Figure S1.8 LCM clusters from CNV Clone B primary cancer in vessel.

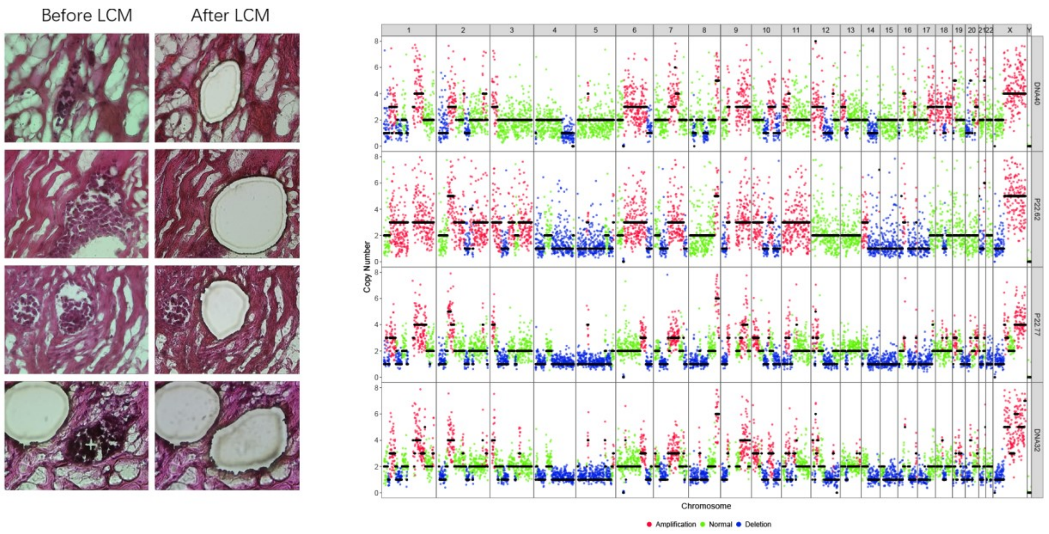


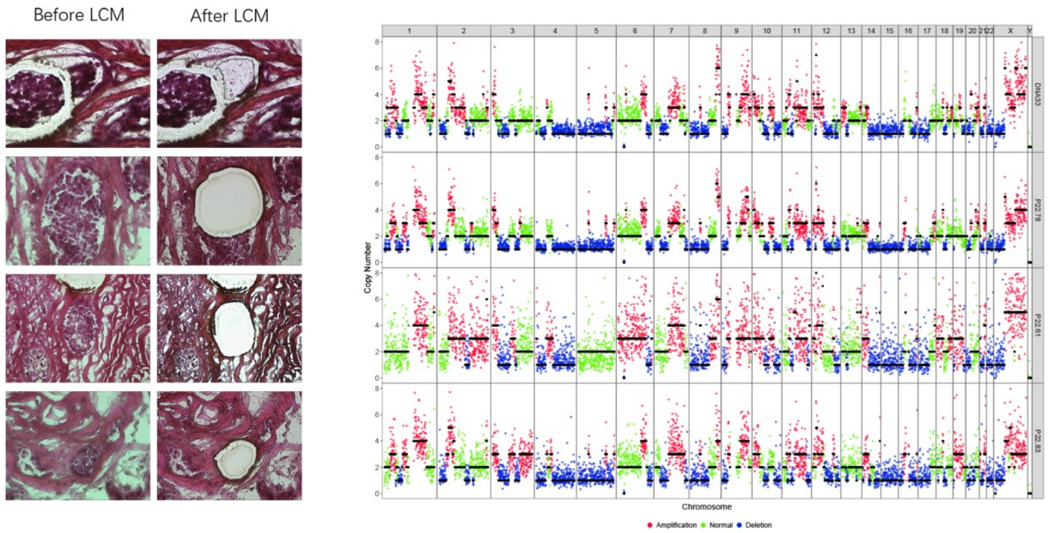


Copy Number Abbreviations for Clone B Primary Cancer in Vessel

Figure S1.9 LCM clusters from CNV Clone B primary cancer in vessel.

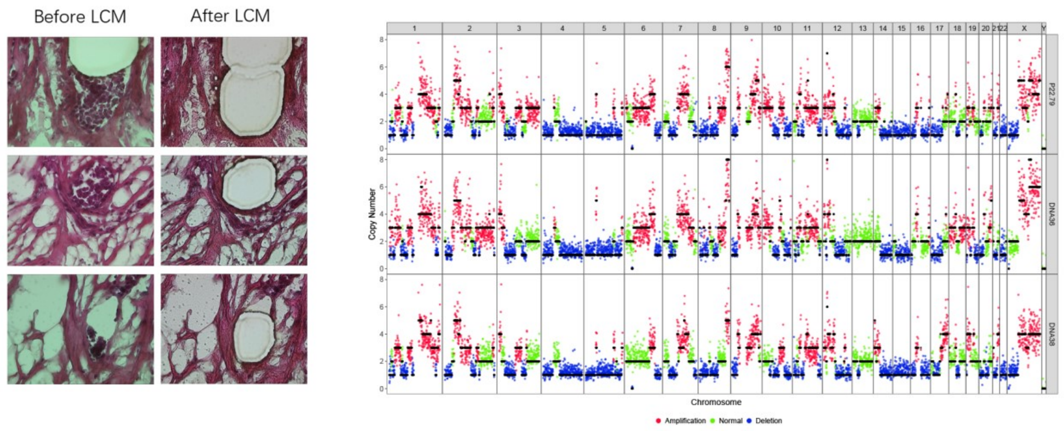


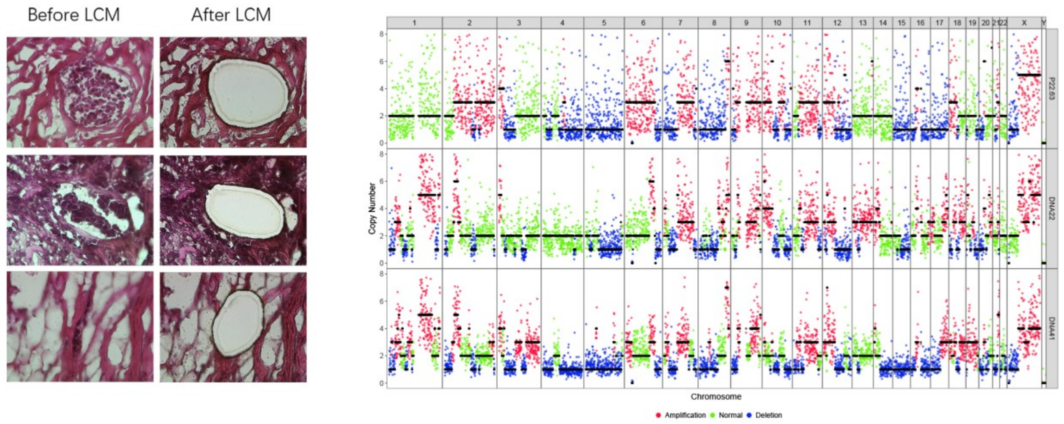


Copy Number Abbreviations for Clone B Primary Cancer in Vessel

Figure S1.10 LCM clusters from CNV Clone C lymph node cancer.

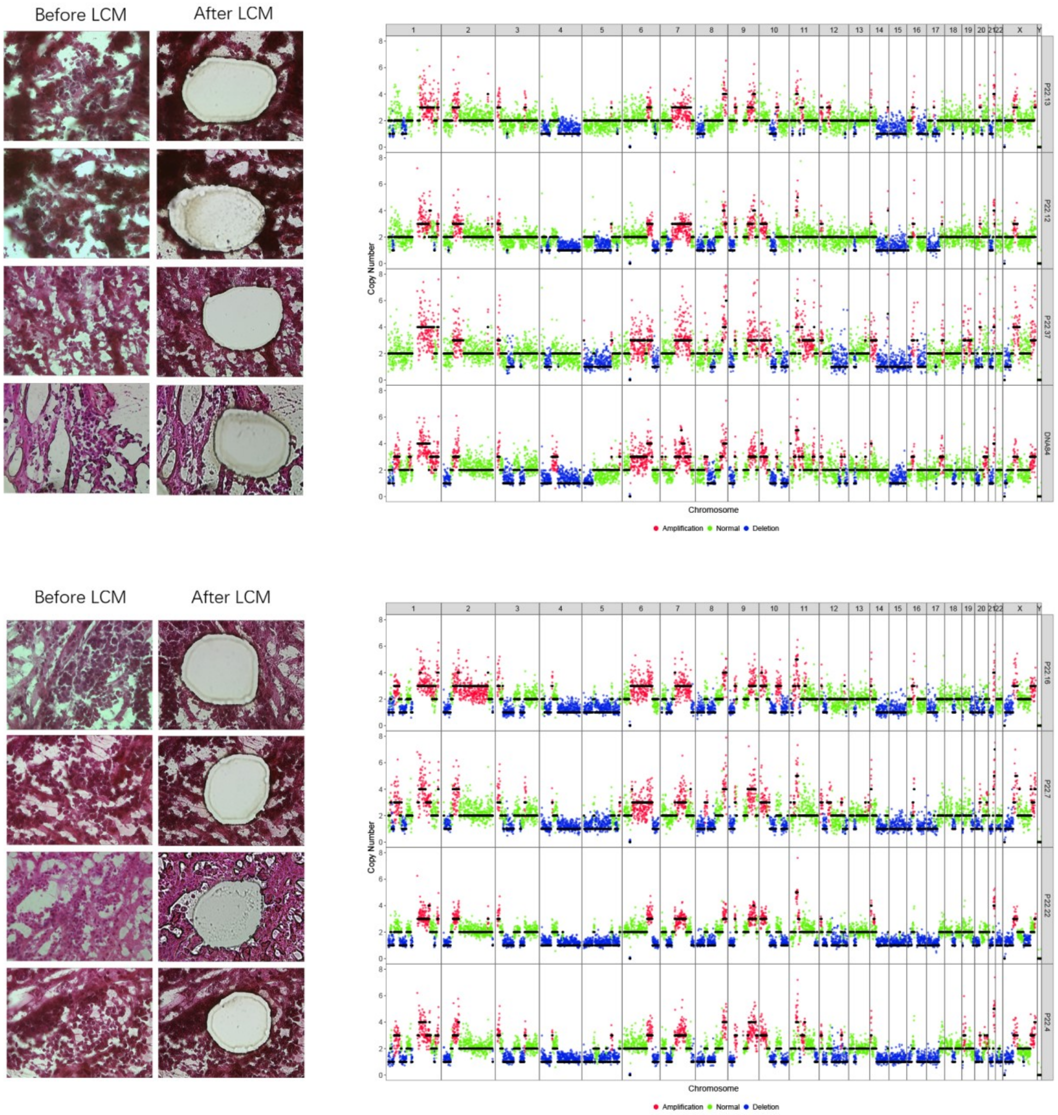


Copy Number Abbreviations for Clone C Lymph node Cancer

Figure S1.11 LCM clusters from CNV Clone C lymph node cancer.

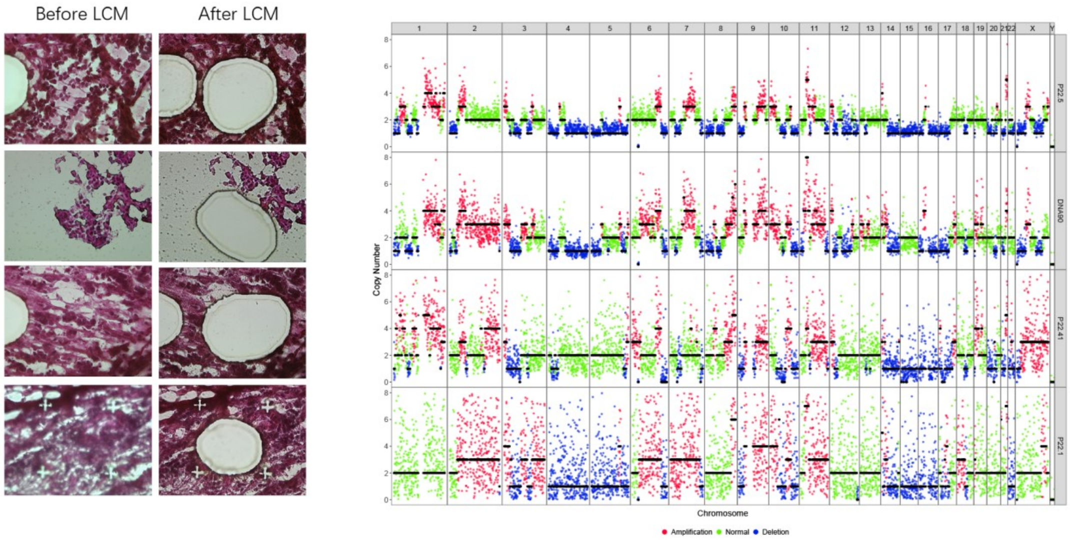


Copy Number Abbreviations for Clone C Lymph node Cancer

FigureS2. SNV heatmap of 1744 non-filtered SNVs (top), and a comparison of the heterozygous and homozygous mutation burden between the three SNV clones (bottom).

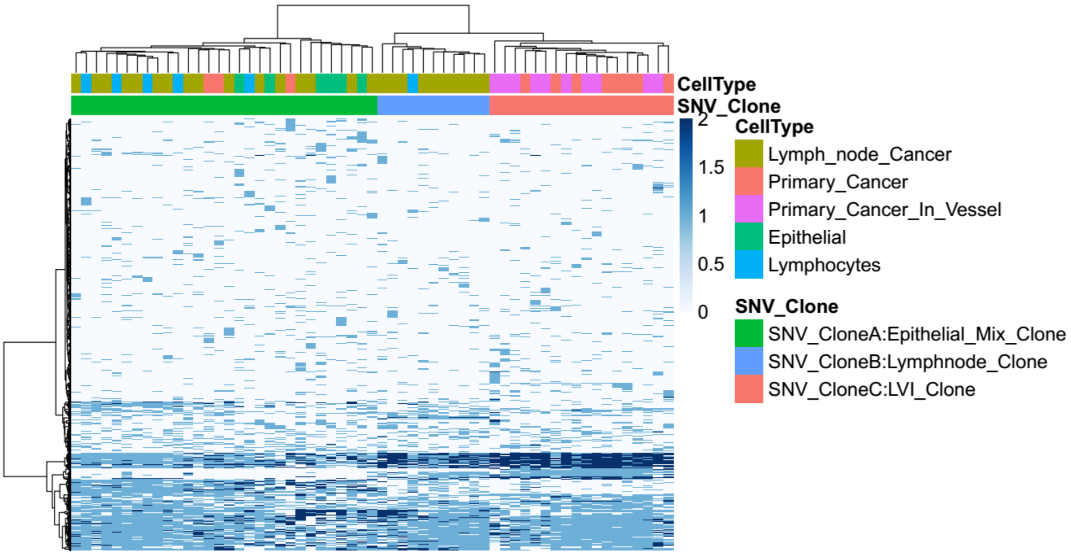


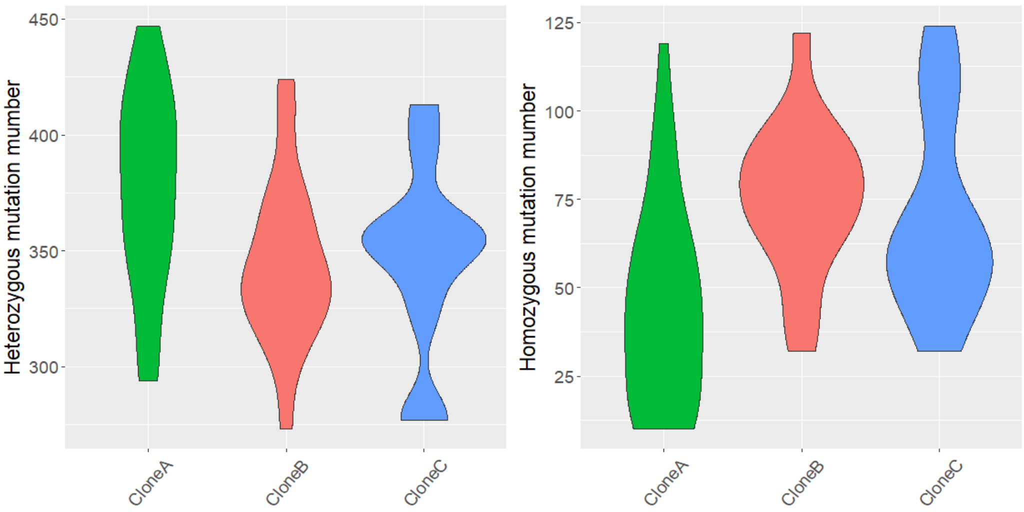


Note: (Top) Heatmap of a total of 1744 SNVs generated by unsupervised hierarchal clustering of 59 cell clusters following WES. Cell clusters are plotted along the x-axis, and SNVs are plotted in genomic order along the y-axis. Color bars (top) represent cell type (yellow, LN; red, primary tumor; pink, primary tumor associated vessels; green, epithelial; blue, lymphocytes), and clone designation (green, mixed clone; blue, LN clone; pink, intravasation clone). Homozygous SNVs are depicted as dark blue, heterozygous SNVs are light blue, and genes with no mutations are white. (Bottom) comparison of heterozygous and homozygous mutation number between SNV clone A (green), clone B (red) and clone C (blue). Mutation number is plotted along the y-axis.

Figure S3. Venn diagrams comparing the number of genes affect by CNV, SNV, and RNA expression changes.

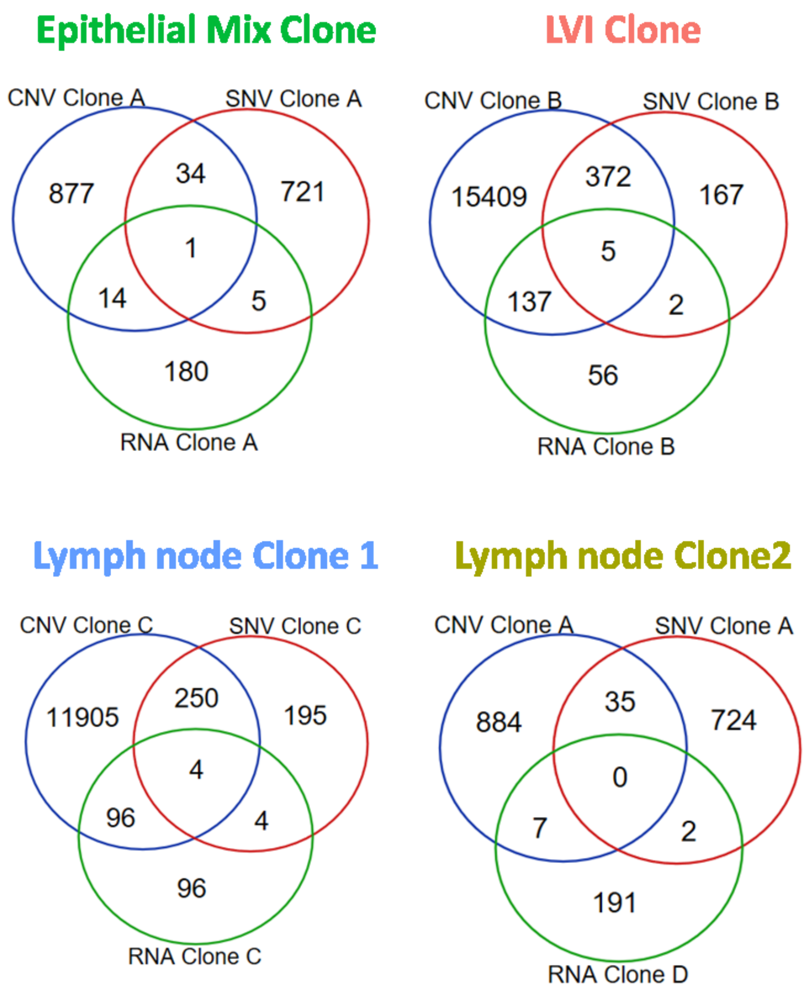


Note: Venn diagrams comparing the number of genes affect by CNV (blue), SNV (red) or transcriptional (green) changes in epithelial, LVI, lymph node 1, or lymph node 2 RNA clone cell clusters.

Figure S4 Venn plot analysis of RNA clones (top), pseudo-time trajectory (middle), and corresponding gene heatmap trajectories of each of the three metastasis paths (bottom).

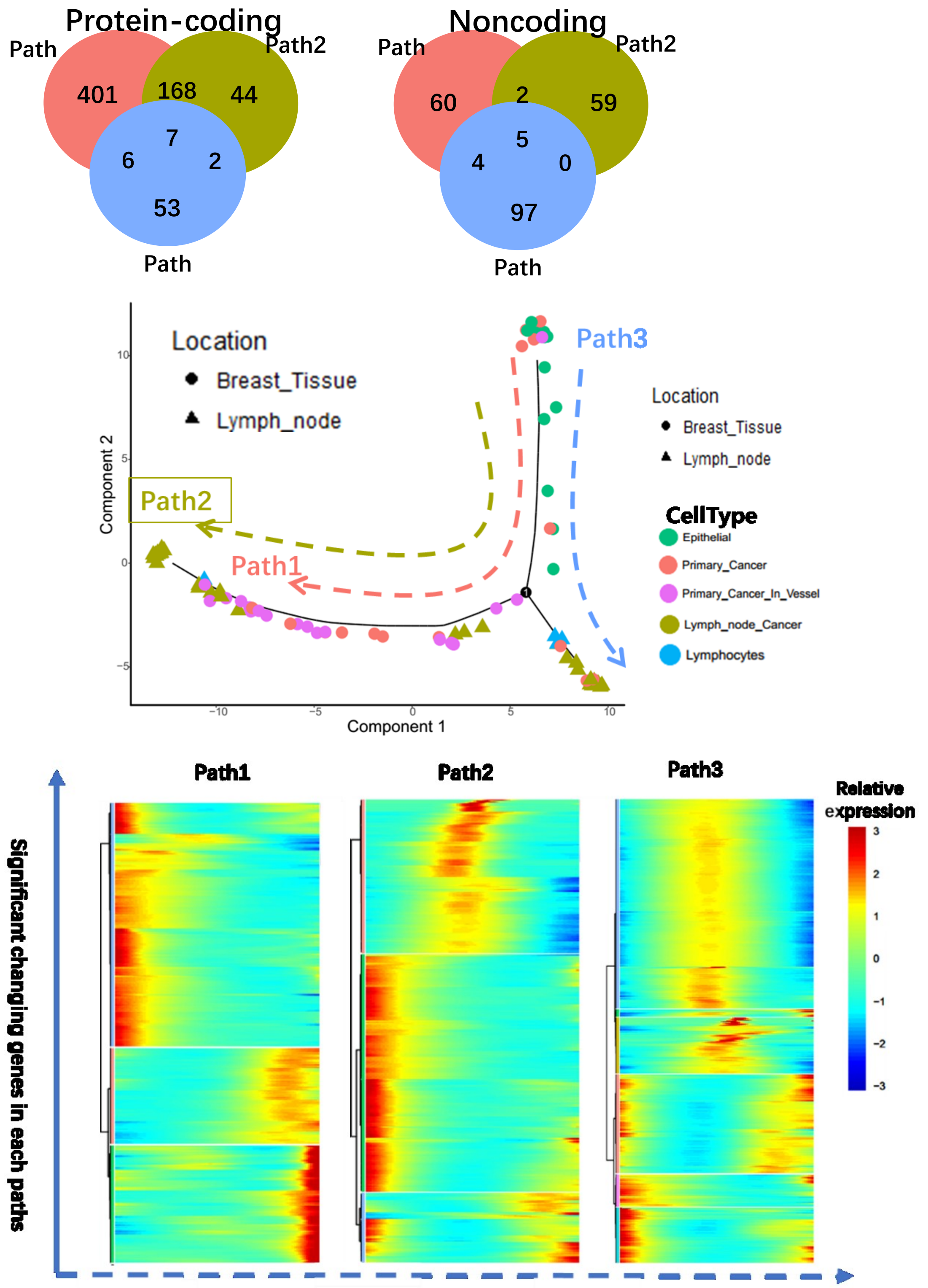


Note: Marker genes were used to generate RNA trajectories for each of three paths and the protein coding and non-coding genes for each trajectory were compared via Veen plot (top). Combined trajectories of each RNA clone as defined by the Monocle 2.0 algorithm (middle). Individual cell clusters are defined by cell type and tissue location. Each marker represents an individual cell cluster isolated from either breast tissue (●) or lymph node tissue (▲). Markers are colored according to cell type (yellow, LN; orange, primary tumor; pink, primary tumor associated lymphatic vessels; green, epithelial cells; blue, lymphocytes). (Bottom) Heatmaps depicting the relative expression of each gene along with pseudo-time trajectories calculated by Monocle2. Pseudo-time is plotted along the x-axis, and the top 800 significant changing genes are plotted along the y-axis.

Figure S5. Enriched biological process networks for the three metastatic paths.

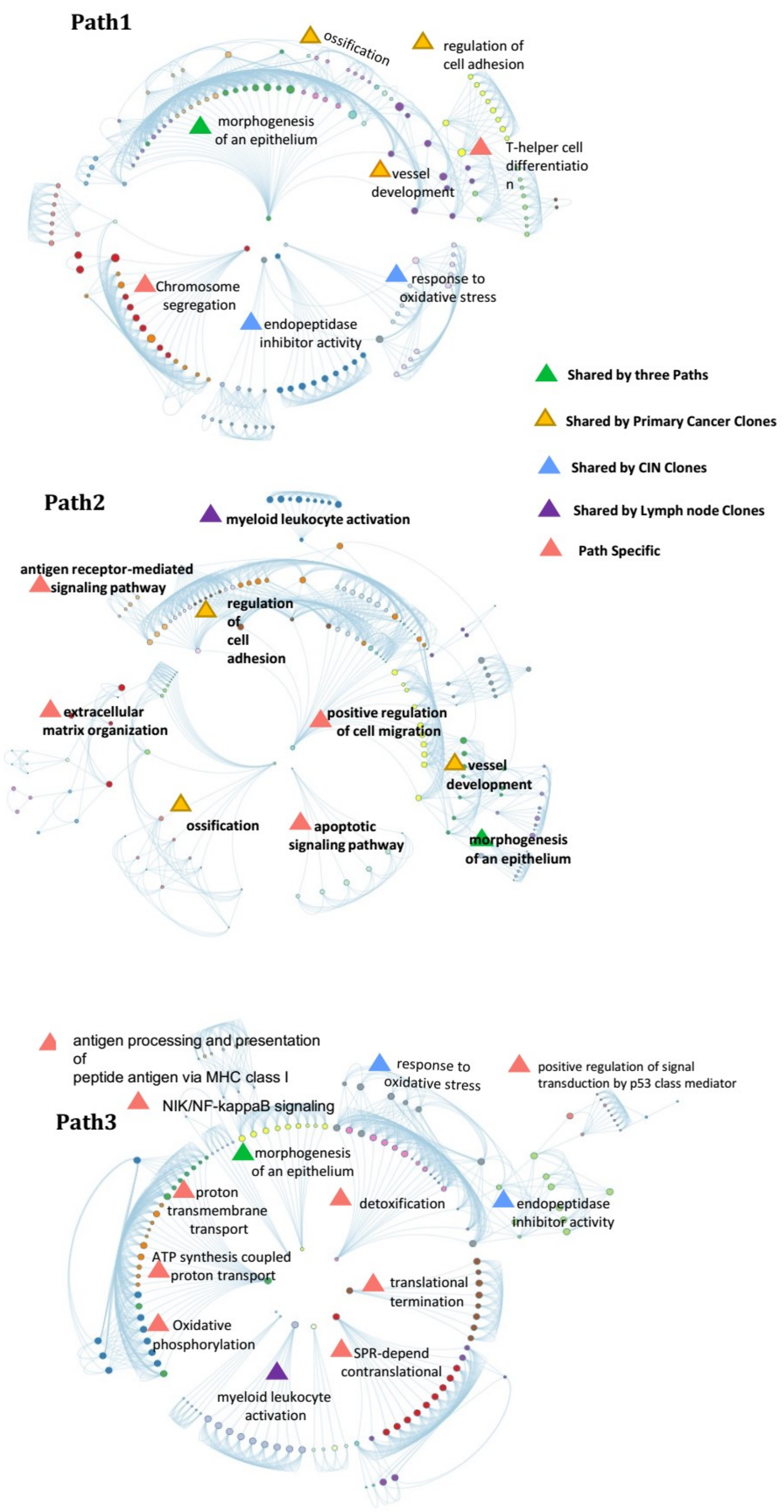


Note: A subset of enriched terms have been selected and rendered as a network plot, where terms with a similarity > 0.3 are connected by edges. We selected the terms with the best p-values from each of the 20 clusters, with the constraint that there be no more than 15 terms per cluster and no more than 250 terms in total with Metascape default setting. The network is visualized using Cytoscape with a radial layout of significantly enriched biological processes, where each node represents an enriched term and is colored first by its cluster ID. Nodes represent individual biological processes and are grouped according to biological pathway Membership. Edges denote known associations between biological processes. Biological pathway Members are further defined as: 1) shared by three paths (green); 2) shared by CIN clones (blue), 3) shared by primary cancer clones (yellow); 4) shared by lymph node clones (purple); and 5) path specific (red).
